# AMPK Drives Both Glycolytic and Oxidative Metabolism in T Cells During Graft-versus-host Disease

**DOI:** 10.1101/2023.06.12.544686

**Authors:** Archana Ramgopal, Erica L Braverman, Lee-Kai Sun, Darlene Monlish, Christopher Wittmann, Manda J. Ramsey, Richard Caitley, William Hawse, Craig A. Byersdorfer

**Author notes:** Corresponding author: Dr. Craig A. Byersdorfer, Division of Blood and Marrow Transplant and Cellular Therapies, Rangos Research Building, 4401 Penn Ave, Pittsburgh, PA 15224. A.R. and E.L.B. contributed equally to this work.

## Abstract

Allogeneic T cells reprogram their metabolism during acute graft-versus-host disease (GVHD) in a process reliant on the cellular energy sensor AMP-activated protein kinase (AMPK). Deletion of AMPK in donor T cells limits GVHD but still preserves homeostatic reconstitution and graft-versus-leukemia (GVL) effects. In the current studies, murine T cells lacking AMPK decreased oxidative metabolism at early timepoints post-transplant and were also unable to mediate a compensatory increase in glycolysis following inhibition of the electron transport chain. Human T cells lacking AMPK gave similar results, with glycolytic compensation impaired both *in vitro* and following expansion *in vivo* in a modified model of GVHD. Immunoprecipitation of proteins from day 7 allogeneic T cells, using an antibody specific to phosphorylated AMPK targets, recovered lower levels of multiple glycolysis-related proteins including the glycolytic enzymes aldolase, enolase, pyruvate kinase M (PKM), and glyceraldehyde 3-phosphate dehydrogenase (GAPDH). Functionally, murine T cells lacking AMPK exhibited impaired aldolase activity following anti-CD3/CD28 stimulation and a decrease in GAPDH activity on day 7 post-transplant. Importantly, these changes in glycolysis correlated with an impaired ability of AMPK KO T cells to produce significant amounts of interferon gamma (IFNγ) upon antigenic re-stimulation. Together these data highlight a significant role for AMPK in controlling oxidative and glycolytic metabolism in both murine and human T cells during GVHD and endorse further study of AMPK inhibition as a potential target for future clinical therapies.

**KEY POINTS:** AMPK plays a key role in driving both and oxidative and glycolytic metabolism in T cells during graft-versus-host disease (GVHD)

Absence of AMPK simultaneously impairs both glycolytic enzyme activity, most notably by aldolase, and interferon gamma (IFNγ) production

## INTRODUCTION

Allogeneic stem cell transplantation (alloHSCT) is a curative treatment for a variety of hematological disorders including relapsed or refractory leukemia, bone marrow failure, and primary immunodeficiencies. In the case of refractory leukemias, alloHSCT provides the added benefit of a graft-vs-leukemia (GVL) response. However, while alloHSCT can be a life-saving procedure, it carries the risk of acute graft-versus-host disease (aGVHD), where donor T cells attack vital organ systems and destroy tissues in the liver, gastrointestinal tract, and skin ^1^. In many cases, the first-line treatment for aGVHD continues to be corticosteroids, although recent additions to the GVHD armamentarium have included agents such as the janus kinase inhibitor Ruxolitinib ^2^. However, in addition to the undesirable consequences of reduced GVL efficacy and increased risks of infection common to most immunosuppressive medications ^3^, many agents also carry drug-specific side effects including hypertension, hyperglycemia, and psychiatric concerns for steroids, nephrotoxicity for calcineurin inhibitors ^4^, and marrow suppression in the case of Ruxolitinib ^5^. Further, even first line therapies continue to show limited utility, with long-standing GVHD remission rates approximating 55% in many patients ^6^. Thus, treatments to specifically target GHVD-causing T cells, while still maximizing physiologic immunity, are still urgently needed ^7^.

Recent literature has elucidated a strong connection between metabolism and subsequent T cell function, where manipulating metabolic decision-making in T cells can impact their cytotoxicity and survival ^8–10^. Alloreactive T cells experience significant metabolic demands in the early post-transplant period, with day 7 alloreactive T cells simultaneously increasing both oxygen consumption and extracellular acidification, a proxy for glycolysis ^11^. This finding is supported by reports that increased aerobic glycolysis, GLUT1 expression, fat oxidation, and glutaminolysis are necessary for alloreactive T cell function and survival ^12–16^. In contrast, processes like lymphopenic or homeostatic reconstitution are often less energy intensive for the donor T cells ^14, 17^ and support the idea that alloreactive T cells specifically adapt their metabolism secondary to the robust demands of responding to nearly ubiquitous antigen presentation in a continuous fashion. This paradigm also highlights the possibility of targeting a cell’s adaptation potential, rather than a specific metabolic pathway, to therapeutically discriminate between alloreactive and non-alloreactive T cells, or between alloreactive T cells with high versus low levels of activation. In this scenario, targeting metabolic adaptation would selectively impair only those cells with increased energy needs (alloreactive T cells) while sparing cells engaged in lower energy tasks (homeostatic reconstitution). One adaptor known to play a robust role in metabolic decision-making is the cellular energy sensor AMP-activated protein kinase (AMPK).

AMPK is a heterotrimeric protein complex composed of α, β and γ subunits. The α subunit contains the kinase activity, the β subunit contributes a structural role and some substrate recognition, and the γ subunit plays a regulatory role. When energy stores are low, a rise in the AMP/ATP ratio activates AMPK through phosphorylation of Thr172 on the α subunit, which then drives AMPK-mediated phosphorylation of downstream targets to promote catabolic energy generation while blocking anabolic growth^18, 19^. The γ subunit can further potentiate AMPK activation by protecting the phosphorylated status of the Thr172 residue^20^ and thereby prolong downstream phosphorylation events. We have previously shown that AMPK activity increases in allogeneic T cells and that deletion of AMPK in T cells prior to transplantation reduces aGVHD severity while still preserving GVL effects^14^. These findings are supported by a role for AMPK in T cell’s adaptation to multiple other stressful *in vivo* environments^21–23^. However, despite a recognized decrease in total AMPK KO T cell numbers early post-transplant, as well as disadvantage to AMPK KO T cells during competitive transplantation, the *in vivo* metabolic consequences of deleting AMPK have not been well-characterized. In addition, it is unknown if human T cells depend on AMPK during GVHD similar to what is seen in murine T cells. Finally, it is unclear whether or how changes in T cell metabolism following AMPK deletion might relate to the differential ability of cells to execute GVHD versus GVL responses.

In these studies, we demonstrate that AMPK is necessary for maximal oxygen consumption and spare respiratory capacity (SRC) in both murine and human T cells. AMPK deficient T cells also lack a compensatory increase in glycolysis, particularly following electron transport chain inhibition. This decreased glycolytic response occurs concomitant with a decrease in the per cell expression of IFNγ, a critical player in GVHD pathogenesis^24^, and these results are consistent with the well-known link between IFNγ production and metabolic pathway utilization^25, 26^. Together, these data build a paradigm where AMPK increases oxidative and glycolytic metabolism in alloreactive T cells in response to metabolic stress and highlight the potential clinical utility in targeting this metabolic decision-making in order to curtail T cell-mediated inflammatory responses, with significant therapeutic implications for the prevention and treatment of aGVHD.

## METHODS

### Mice

C57BL/6 (B6, H2^b^), B6 x DBA2 F1 (B6D2F1), CD45.1 (B6.SJL-Ptprca^Pepcb/Boy^J), CD4cre (B6.Cg-Tg(Cd4-cre)^1Cwi/Bflu^J), NSG (NOD.*Cg-Prkdc^scid^Il2rgt^m1Wjl^*/SzJ), and Thy1.1 (B6.PL-Thy1^a^/CyJ) mice were purchased from Jackson Laboratories. AMPKα1^fl/fl^α2^fl/fl^ mice were a kind gift from Sean Morrison^27^ and backcrossed to C57Bl/6 mice for >6 generations. Male and female mice were used interchangeably. Recipient animals were 8-12 weeks old, and donor animals 8-16 weeks old, at the time of transplantation. All animals were housed in a specific pathogen-free facility.

### Bone marrow transplantation

Unless otherwise stated, B6D2F1 mice were conditioned with 1100 cGy total body irradiation (TBI) in a split dose from an X-ray source (X-rad 320, Precision X-ray, N. Branford, CT). This was followed by intravenous infusion of 5×10^6^ T-cell-depleted (TCD) B6 BM cells and 2×10^6^ CD45.1^+^ B6 T cells which had been enriched via CD90.2 positive selection (Miltenyi). In xenogeneic GVHD experiments, NSG mice were irradiated with 160 cGy followed by administration of variable numbers of expanded human T cells and 1×10^6^ recently thawed autologous non-T cell APCs. Transplanted mice were housed in sterile microisolator cages on autoclaved water and irradiated chow, and humanely euthanized for moribund behavior or weight loss >30% from baseline.

### Seahorse Analysis

The Seahorse XF Cell Mito Stress Test Kit (Agilent, Santa Clara, CA; Catalog #103015-100) was run on a Seahorse XFe96 Bioanalyzer (Agilent) to determine basal and maximal rates of oxygen consumption (OCR), SRC, and extracellular acidification (ECAR). T cells were plated in assay media (XF Base media (Agilent) with glucose (25mM), sodium pyruvate (2 mM) and L-glutamine (4 mM) (Gibco), pH 7.4 at 37 °C) on a Seahorse cell culture plate coated with Cell-Tak (Corning) at 1×10^5^ cells/well. After adherence and equilibration, basal ECAR and OCR readings were taken for 30 min. Cells were then stimulated with oligomycin (2 µM), carbonyl cyanide 4-(trifluoromethoxy) phenylhydrazone (FCCP, 1 µM), and rotenone/antimycin A (0.5 µM) to obtain maximal respiratory and control values. Assay parameters: 3 min mix, no wait, 3 min measurement, 3 cycles repeated at baseline and after each injection. SRC was calculated as the difference between basal and maximal OCR values obtained after FCCP uncoupling. The XF Mito Stress Test report generator and Agilent Seahorse analytics were used to calculate parameters from Wave software (Agilent, Version 2.6.1.53).

### CRISPR/Cas9 gene editing

Primary human T cells were isolated from a healthy human buffy coat using the Miltenyi Pan human T Cell Isolation Kit for human cells, and frozen. For knockout experiments, sgRNA targeting the AMPKα1 gene (ATGTGATGGGATCTTCTATA) was obtained from Integrated DNA Technologies (IDT), T cells were thawed, rested for two hours, and subsequently stimulated with CD3/CD28 Dynabeads for 48-72 hours. Cells were then removed from the Dynabeads, counted, and rested for an additional 2-24 hours. 4×10^5^ cells were electroporated following the IDT-published protocol (https://www.idtdna.com/pages/support/guides-and-protocols) using the Neon transfection system (Thermo Fisher) with electroporation enhancer. Electroporator settings were 3x 1600V pulses with a 10ms pulse width using the 10 uL pipette. Following electroporation, cells rested for four hours at 37 degrees in antibiotic-free Dulbecco’s Modified Eagle Medium media, followed by plating in a 96-well plate in AIM V media with 5% immune cell serum replacement (SR) and 100 IU of human IL-2. Cells were given fresh media with IL-2 (100 IU) every 2 days and replated every 4 days at 0.5×10^6^ T cells/ml.

### Detection of Indels via TIDE analysis

To calculate the degree of insertions and deletions, 1×105 cells were removed four days after electroporation and DNA isolated using the DNeasy kit (Qiagen) according to manufacturer’s instructions. PCR products were generated using forward (5’-TTGGCCCTATTGTCAGATCCC-3’) and reverse (5’-AGTGAAAGCCCTCCCTCTTAC-3’) primers with an annealing temperature of 60 C for 45 seconds and PCR products purified via the QIA Quick PCR Purification Kit (Qiagen). DNA concentration was measured on the Nanodrop, and samples were run on an agarose gel to ensure proper band size. Quality-controlled PCR samples were sent for Sanger sequencing to GeneWiz and gene editing analyzed in comparison to the mock group using the tracking of indels by decomposition (TIDE) algorithm^28^.

### Protein Isolation and Immunoblot

1.5×10^5^ human or murine T cells were washed once in PBS and re-suspended in 10% trichloroacetic acid (TCA). Cell lysates were centrifuged at 16,000xg at 4C for ten minutes, washed twice in ice cold acetone, resuspended in solubilization buffer (9M Urea containing 1% DTT and 2% Triton X and NuPAGE lithium dodecyl sulfate sample buffer 4X (Invitrogen) at a 3:1 ratio), and heated to 70C for 10 minutes^29^. Electrophoresis was performed on NuPAGE 4-12% Bis-Tris Protein Gels (Invitrogen) at 135V and proteins transferred to Invitrolon^TM^ 0.45µm PVDF membranes (Invitrogen) at 30V for one hour on ice. Membranes were blocked in Tris Buffered Saline with 2% Triton (TBS-T) containing 5% nonfat milk and immunoblotting performed according to the Cell Signaling Technologies Western Blot Protocol. Blots were stripped for 30 minutes (1% SDS, 25 mM glycine, pH 2.0) prior to re-probing. Antibodies used for immunoblotting are listed in Table S2. Blots were developed with Super Signal West Femto chemiluminescence reagents (Thermo Fisher Scientific), detected by CL-X Posure Film (Thermo Scientific), and scanned in grayscale using an Epson V600 scanner. Images were inverted using ImageJ Software (version 1.47T) and densitometry quantitated in an area encompassing the largest band, followed by quantitation of all subsequent bands using the same 2-dimensional area.

### Flow Cytometry

Cells were washed in PBS with 2% fetal bovine serum (FBS) before staining with antibodies at 1:100 dilution for 30 minutes. For detection of intracellular cytokines, 3×10^5^ transplanted cells were recovered from the spleens of recipient mice and mixed with 3×10^5^ fresh F1 splenocytes, followed by culturing for 6 hours in the presence of 1mM Brefeldin A. Cells were subsequently stained for extracellular markers, washed twice, and fixed per manufacturer’s instructions (Fix/Perm kit, Invitrogen Cat #:88-8824-00). Intracellular cytokine staining utilized a 1:100 dilution of antibody in permeabilization buffer for 30 minutes at room temperature, followed by two washes, and subsequent analysis. Flow data was captured on a BD Fortessa analyzer (BD Biosciences) and evaluated using FlowJo software (version 10.1, Tree Star). Intracellular staining of *in vitro* activated cells occurred similarly, with the exception that cells were re-stimulated for 6 hours in the presence of Brefeldin A on anti-CD3, anti-CD28 antibody coated plates.

### ELISA and cytokine multiplex analysis

Levels of murine cytokines in transplanted mice were assessed using the LEGENDplex™ Murine Inflammation panel (13-plex) in a V-bottom plate per manufacturer’s instructions (Cat. No: 740446, BioLegend, San Diego, CA, USA). LEGENDPlex data were acquired on a BD Fortessa analyzer (BD Biosciences, Franklin Lakes, NJ, USA) and assessed using FlowJo software (version 10.7, Tree Star). Human IFNγ levels were detected via enzyme linked immunosorbent assay (ELISA) using the Human IFNγ Quantikine ELISA kit (R&D Systems) per manufacturer’s instructions. Serum was analyzed at a 1:4 dilution.

### Immunoprecipitation of AMPK substrates and mass spectrometric analysis

A total of 1×10^6^ T cells were collected and lysed with a buffer containing 25 mM Tris–HCl, pH 7.4, 150 mM NaCl, 1 mM EDTA, 1% Nonidet P-40, 5% glycerol that was supplemented with Roche Complete C protease and PhosSTOP phosphatase inhibitors. Lysates were sonicated and centrifuged to remove insoluble debris, and the remaining lysate was incubated with Phospho-AMPK Substrate Motif mAb mix (Cell Signaling Technologies; catalog No. 5759) for 12 h at 4 °C. Antibody complexes were incubated with protein A beads (Cell Signaling Technologies; catalog No. #9863) for 1 h at room temperature. Beads were washed twice with lysis buffer and proteins were eluted from the beads with 8 M urea (U5128; Sigma) and 0.1 M Tris–HCl at pH 8.5. The filter-aided sample preparation method was used to generate tryptic peptides and desalted using C18 spin column ^30^. Peptides were suspended in 0.1% formic acid and resolved with liquid chromatography tandem mass spectrometry using a system composed of a Waters nanoACQUITY HPLC in-line with either an LTQ/OrbitrapVelos Elite hybrid mass spectrometer (Thermo Fisher). Solvent A (0.1% formic acid in HPLC-grade water) and solvent B (0.1% formic acid in 100% acetonitrile) were used as the mobile phase. Peptides were then eluted onto a capillary column (75 μm inner diameter × 360 μm outer diameter × 15 cm long; Polymicro Technologies) 5-μm particle size, 125 pore size C-18 silica-bonded stationary phase (Phenomenex) and resolved using a 100-min gradient at the flow rate of 0.2 μl min−1 (3–33% B for 90 min, 33–80% B for 2 min, constant at 80% B for 6 min, and then 80–0% B for 2 min to equilibrate the column). Data were collected in positive ionization mode. PEAKS9 software was used to sequence and identify peptides in each sample using a decoy search at a 1% false discovery rate using the UniProt murine database. Label-free quantitation was performed using the quantitative module in the PEAKS9 software.

### Glycolytic enzyme assays

Aldolase A and GAPDH enzymatic assays were run according to manufacturer’s instructions (Biovision-Abcam) on cell lysates from 2.5×10^5^ T cells per well. To detect aldolase activity, T cells were collected after 3 days of stimulation with anti-CD3/anti-CD28 antibodies. To measure GAPDH activity, T cells were positively selected for CD45.1 from day 7 recipient spleens, followed by cell lysis and detection of enzyme activity.

### Mixed leukocyte reactions

Murine T cells were purified from unmanipulated donors via column-based positive selection for CD90.2 (Miltenyi), labeled with CellTrace Violet, and plated at 3×10^5^/well with 3×10^5^ splenocytes from B6D2F1 animals. MLRs were performed on 96-well flat plates in Dulbecco’s Modified Eagle’s Media (Gibco) supplemented with 10% FBS, L-glutamine, nonessential amino acids, sodium pyruvate, and penicillin/streptomycin (all from Life Technologies). Cultures were assessed at 72 hours for cell division status and subsequently analyzed by flow cytometry for intracellular cytokine production.

### Statistics

Graphing and statistical analysis was performed using GraphPad Prism for Windows (version 9.3.0, San Diego, CA; www.graphpad.com). Unpaired two-tailed Student t test was used to determine statistical significance. Unless noted otherwise, data are displayed as mean ± standard deviation. In all cases, *p<0.05, **p<0.01, ***p<0.001, and ****p<0.0001.

## RESULTS

### AMPK KO T cells reduce both oxidative and glycolytic metabolism post-transplant

Previous work has demonstrated that donor T cells lacking AMPK mediate less severe GVHD, but the metabolic underpinnings of this reliance were not previously explored. Given the known role of AMPK as a master regulator of oxidative metabolism^31, 32^, we wished to test the oxidative capacity of AMPK KO T cells from our GVHD model. Wildtype (WT) or AMPK KO T cells were transplanted into B6D2F1 recipients, recovered on day 7 via negative selection from the spleens of recipient animals, and analyzed using the mitochondrial stress kit on the Seahorse metabolic analyzer. Baseline oxygen consumption rates (OCR) were equivalent between WT and AMPK KO groups (**Figure 1A**), but AMPK KO T cells decreased both their maximal OCR and SRC, suggesting that AMPK KO T cells were already operating near their maximal oxidative capacity at baseline, with very little left in reserve (**Figure 1B-C**). We next interrogated glycolysis by measuring rates of extracellular acidification (ECAR). Again, baseline ECAR values were similar between WT and AMPK KO T cells (**Figure 1D**). However, AMPK KO T cells failed to increase glycolysis upon electron transport chain (ETC) inhibition with oligomycin (**Figure 1D-E**), resulting in diminished maximal ECAR values in AMPK KO T cells (**Fig. 1F**). Together, these data demonstrate a role for AMPK in enhancing both oxidative and glycolytic metabolism in GVHD T cells, particularly in response to an acute metabolic stress (e.g. oligomycin), suggesting an important function for AMPK in facilitating early metabolic adaptation of T cells to the allogeneic environment.

**Figure 1.**
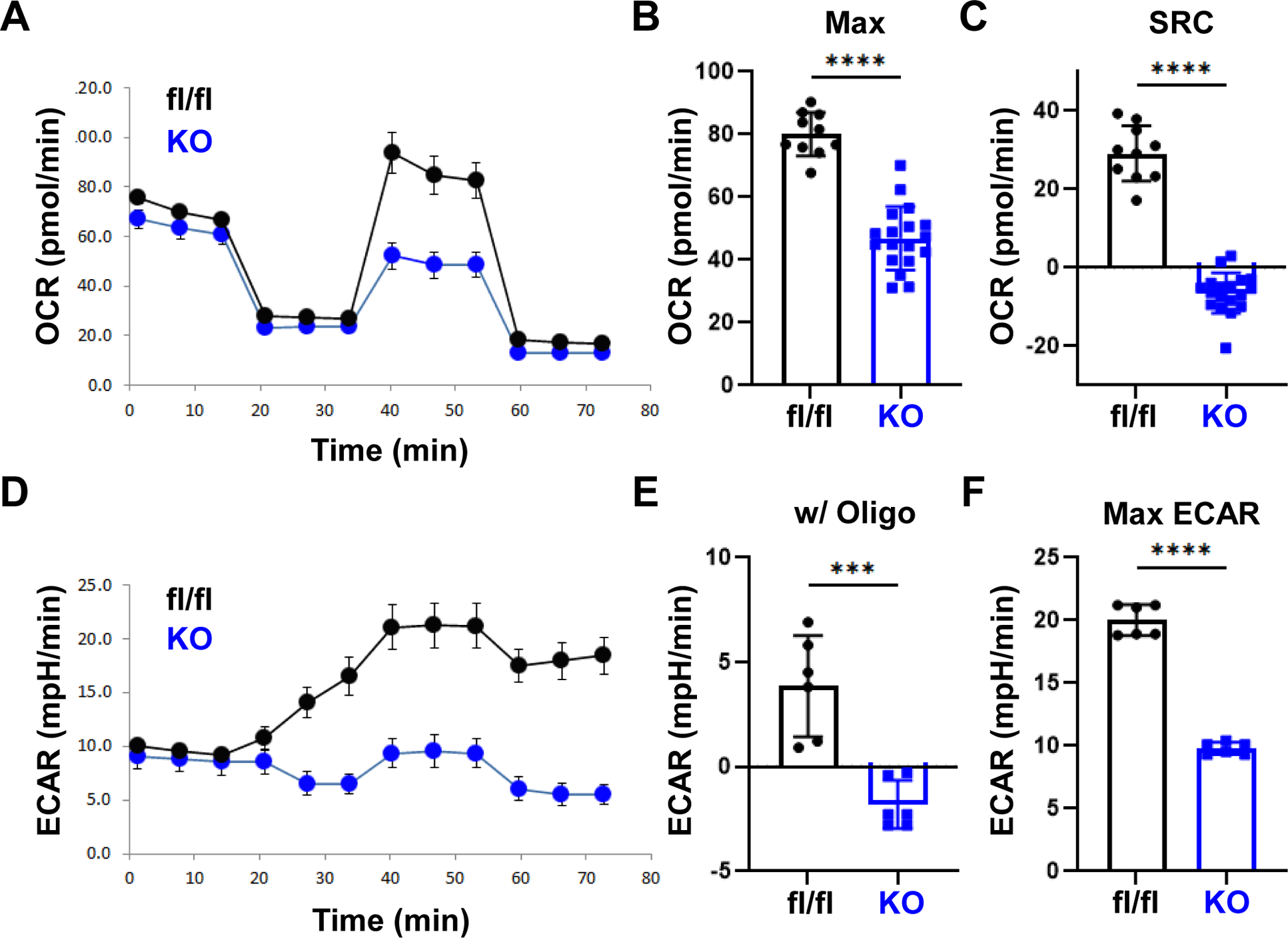
AMPK KO T cells reduce both oxidative and glycolytic metabolism. 2×10^6^ CD45.1^+^ WT or AMPK KO (blue) T cells and 5×10^6^ T-cell depleted (TCD) B6 bone marrow (BM) cells were transplanted into irradiated allogeneic (B6D2F1) recipients. On day 7 post-transplant, donor T cells were purified by negative selection over a magnetic column, placed into the Seahorse metabolic analyzer, and metabolism interrogated using the mitochondrial stress kit. Oxygen consumption rates (OCR) (**A**), including both maximal OCR (**B**) and spare respiratory capacity (**C**), were measured simultaneously with extracellular rates of acidification (ECAR), as shown in (**D**). Response to oligomycin (w/ oligo) was calculated by subtracting individual values from the averaged baseline values prior to oligomycin administration (**E**). Maximal ECAR values were simply the highest ECAR values obtained over the course of the analysis (**F**). n = 2 pooled samples per group (3-4 mice in each pool) and plots are representative of two independent experiments. ***p<0.001, ****p<0.0001.

### Human T cells lacking AMPK reduce oxidative and glycolytic metabolism *in vitro*

Given the impaired metabolism in AMPK KO murine T cells, we determined if similar changes occurred in human T cells lacking AMPK, a result which could be leveraged therapeutically. To generate human cells lacking, CD3+ human T cells were purified from the peripheral blood of healthy human donors, electroporated with ribonucleoprotein (RNP) complexes containing the Cas 9 protein and a guide RNA (gRNA) targeting human AMPKα1 and expanded for 7-10 days in media containing recombinant human IL-2. Targeting human AMPKα1 in this manner consistently achieved insertion-deletion efficiencies >90% as quantified by TIDE analysis (**Figure 2A**). Loss of AMPK function was further confirmed by immunoblot, where AMPK-specific phosphorylation of the target protein Unc51-Like Kinase (ULK-1) was reduced by more than 85% compared to WT cells (**Figure 2B**). To evaluate the effects of AMPK deletion on metabolic potential, human T cells lacking AMPK were restimulated overnight in physiological glucose (5.5 mM) and analyzed the following day on the Seahorse metabolic analyzer. Similar to murine T cells, human T cells lacking AMPK decreased both maximal OCR (**Figure 2C-D**) and SRC (**Figure 2E**) compared to T cells which were electroporated without the gRNA (mock). AMPK deficient human T cells also decreased ECAR values at baseline, which separated further upon inhibition of the ETC (**Figure 2F-H**).

**Figure 2.**
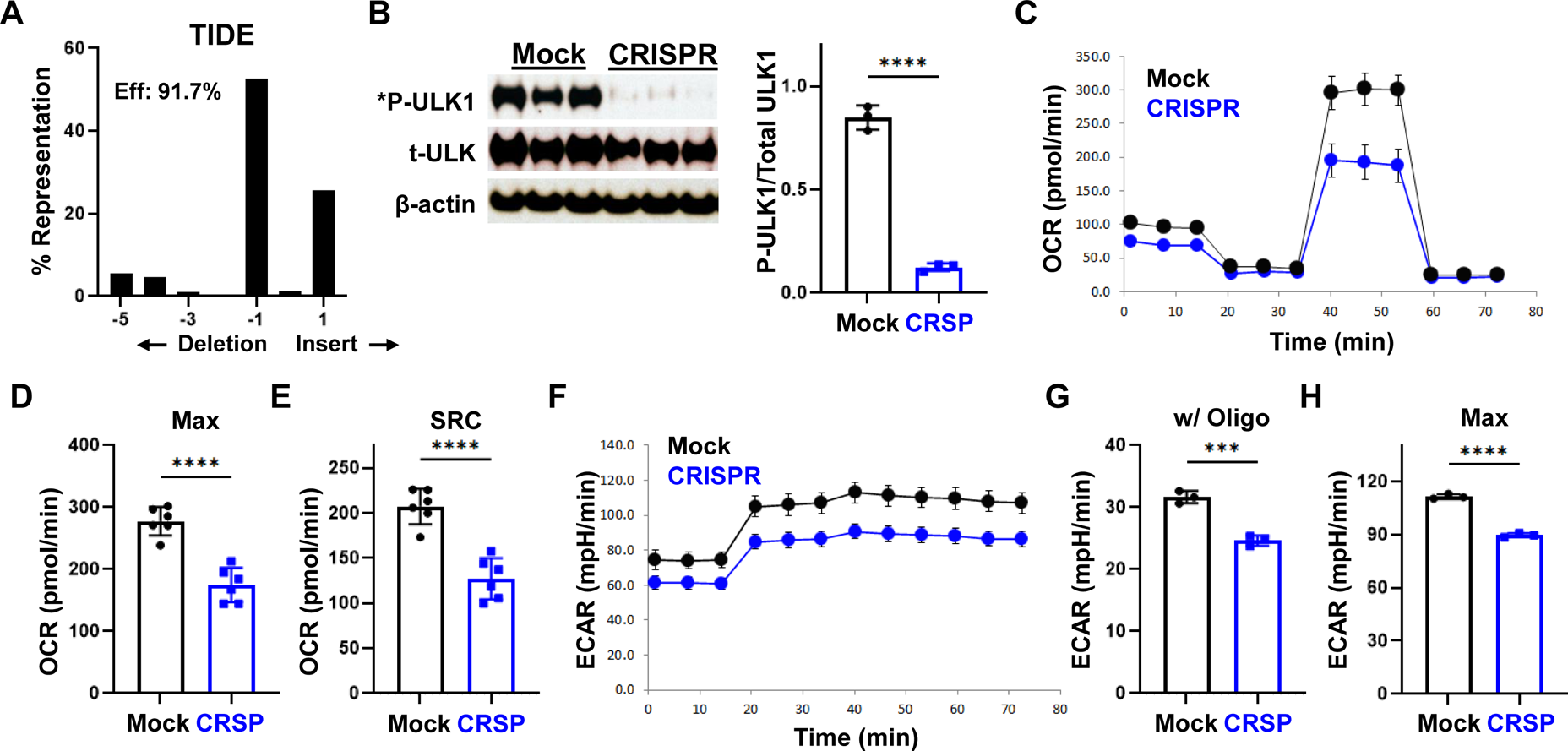
AMPK deficient human T cells decrease oxidative and glycolytic metabolism *in vitro*. CD3+ human T cells were purified from the peripheral blood of healthy human donors, electroporated with ribonucleoprotein (RNP) complexes containing the Cas9 protein and a gRNA targeting human AMPKα1 locus and expanded until day 10 in recombinant human IL-2. *A*, Genomic DNA purified from day 10 T cells was used to amplify a 700 bp fragment covering the gRNA target site, followed by Sanger sequencing and decomposition analysis ^28, 69^. *B*, Protein from 1×10^5^ day 10 T cells was precipitated with TCA followed by immunoblot analysis for specific phosphorylation of ULK-1 on Ser555 by AMPK. *C-H*, Day 10 T cells were stimulated overnight with CD3/CD28 Dynabeads in physiologic glucose (5.5mM) and placed on the Seahorse metabolic analyzer (**C**). Both maximal OCR (**D**) and SRC (**E**) were measured simultaneously with ECAR (**F-H**), including both the response to oligomycin (**G**) and the maximal ECAR values (**H**). Plots in C and F are representative of two, independent donors while data in D-G are the composite analysis from these two donors. ***p<0.001, ****p<0.0001.

### Creating a modified model of xenogeneic GVHD

We next evaluated the metabolism of CRISPR-treated cells in a xenogeneic model of GVHD. To start, we transplanted CRISPR-treated cells into NOD-scid IL2Rgamma^null^ (NSG) mice irradiated with 160 cGy. However, this approach yielded few human T cells, negligible weight loss, and minimal cytokine production by day 25 post-transplant. Noting that previous xenogeneic models used by ourselves^14^ and others injected whole populations of peripheral blood mononuclear cells (PBMCs) as donor cells ^33^, we reasoned that maximal GVHD severity may require non-T cell antigen presenting cells (APCs) in the donor inoculum^34^. Based on this hypothesis, we added 1×10^6^ autologous non-T cell APCs to our day 10 *in vitro*-expanded T cells prior to injection into NSG recipients. Inclusion of 1×10^6^ APCs increased both the percentage and total number of human CD45+CD3+ T cells recovered on day 25 post-transplant (**Figure 3A**). In addition, we could now readily detect elevated levels of human interferon gamma (IFNγ) in the serum of transplanted mice (**Figure 3B**). We next sought to determine if there was a dose-dependent correlation between the number of T cells injected and the quantity of cells recovered on day 25. We also wished to identify an optimal T cell dose for further studies. T cells were diluted in 3-fold dilutions (6×10^6^ down to 0.67×10^6^ T cells/recipient), mixed with 1×10^6^ APC, and transplanted into lightly irradiated recipients. Quantitation of CD45+CD3+ percentages occurred on day 25 post-transplant (**Figure 3C**). Lower starting doses of human T cells yielded proportionally fewer human T cells on day 25 post-transplant (**Figure 3D**), a result which correlated with reduced serum levels of human IFNγ (**Figure 3E**). Changes in donor T cell number also correlated with serum levels of murine CCL-2, which served as an accessory biomarker of disease severity in this modified GVHD model (**Figure 3F**). In total, the standard xenogeneic GVHD model was successfully modified to yield both consistent cell numbers and reliable serum biomarkers by day 25 post-transplant using *in vitro*-expanded human T cells and recently thawed autologous APCs. We ultimately chose 6×10^6^ T cells and 1×10^6^ autologous APCs for subsequent studies.

**Figure 3.**
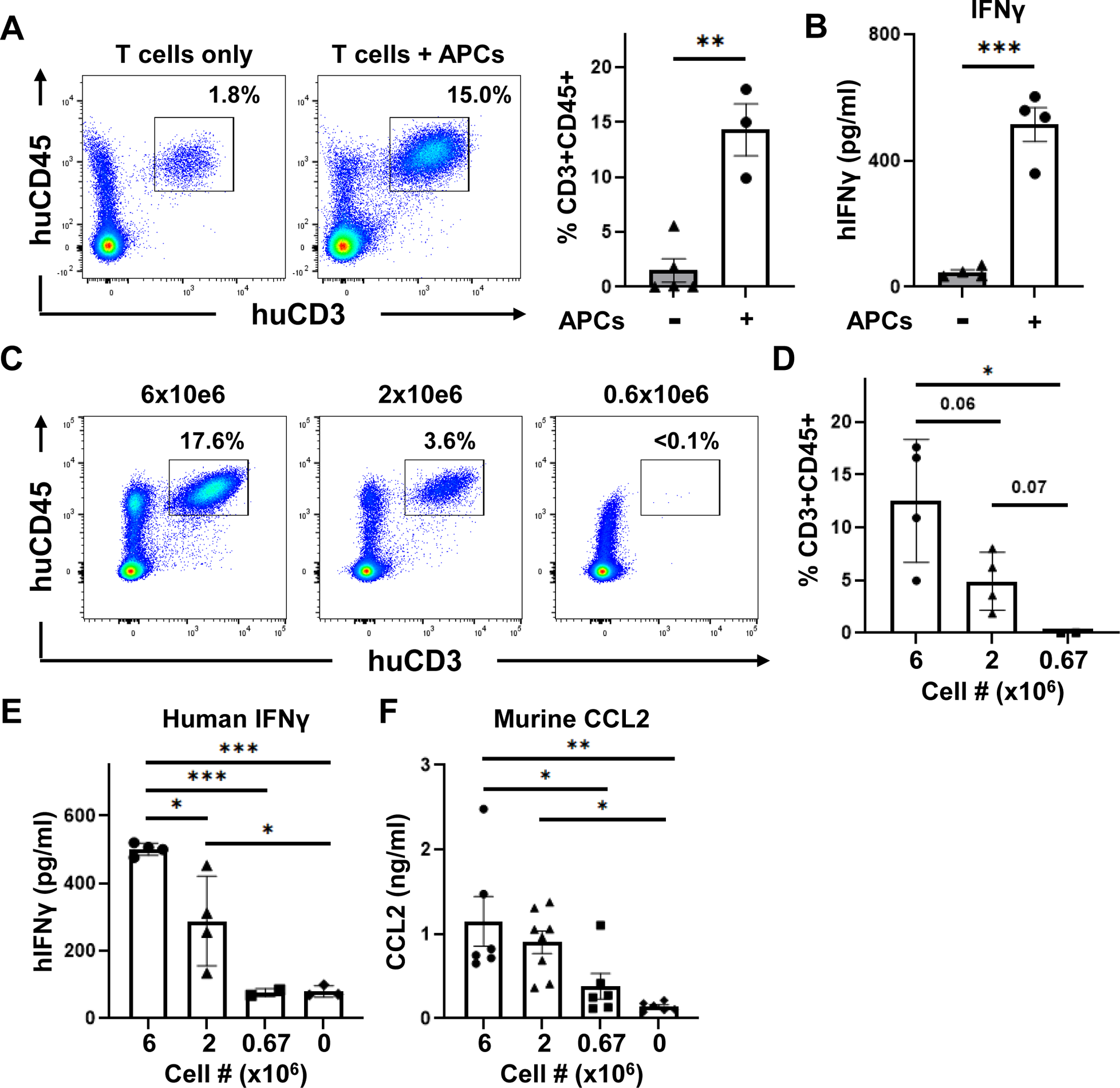
Non-T cell APCs are critical to the development of xenogeneic GVHD. 6×10^6^ human T cells from day 10 of *in vitro* expansion were injected, with or without 1×10^6^ autologous non-T cell APCs, into irradiated NSG recipients. Spleens were recovered on day 25 post-transplant and the percentage of human CD3+CD45+ cells enumerated by flow cytometry (**A**), with serum levels of human IFN-γ measured in these same recipients (**B**). *C-F*, Variable doses of day 10 human T cells were transplanted with 1×10^6^ autologous APCs into irradiated NSG mice and the percentage of human CD3+CD45+ measured in recipient spleens on day 25 post-transplant (**C-D**). Levels of human IFNγ (**E**) and murine CCL2 (**F**) were measured in these same recipients. n=4 mice/group in A-B, with data representative of two independent experiments. C-D are representative data from two additional experiments (n=4 mice/group). E is a single experiment (representative of three total) with n=2-4 mice/group and F is composite data from three individual experiments (n=6-8 mice/group total). *p<0.05, **p<0.01, ***p<0.001.

### Human T cells lacking AMPK decrease glycolytic metabolism *ex vivo*

Following model optimization and determination of the appropriate T cell dose, we sought to characterize AMPK’s role in human T cells using this modified model of xenogeneic GVHD. Human T cells were electroporated with AMPKα1 targeting RNPs, expanded in hIL-2 until day 10, and co-transplanted with 1×10^6^ autologous APCs into lightly irradiated NSG recipients. Importantly, human T cells recovered on day 25 post-transplant continued to remain highly edited, with indel efficiencies above 87% (**Figure 4A**). However, in contrast to AMPK KO T cells in the murine system, human T cells lacking AMPK did not exhibit a decrease in oxidative capacity on day 25 post-transplant (**Figure 4B**) and experienced only minimal changes in baseline ECAR values (**Figure 4C**). However, consistent with day 7 murine T cells, human T cells lacking AMPK continued to manifest an impaired compensatory response to oligomycin administration, with the response to oligomycin at 50% of WT levels (**Figure 4D**) and maximal ECAR values <80% of those for mock T cells (**Figure 4E**). Thus, while diminishing AMPK activity in human T cells did not impact oxidative capacity on day 25, as seen with murine KO T cells on day 7, the lack of a compensatory glycolytic response was consistent in both cell types.

**Figure 4.**
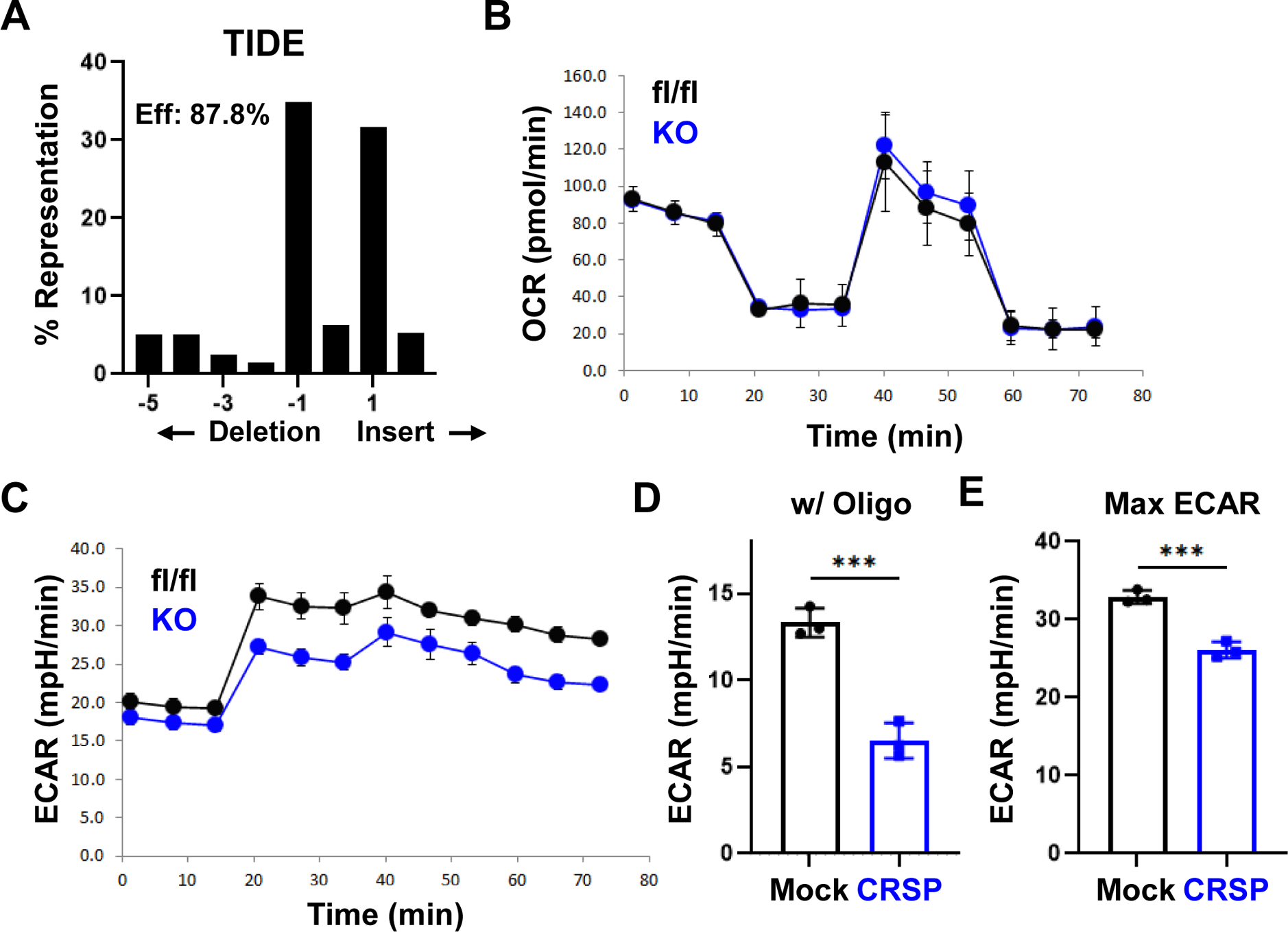
Decreased glycolytic compensation in human T cells lacking AMPK. 6×10^6^ day 10 human T cells were transplanted with 1×10^6^ autologous non-T cells into irradiated NSG recipients. Human CD3+ T cells were recovered from spleens of NSG recipients on day 25 post-transplant and purified via magnetic selection. *A*, Genomic DNA was purified from day 25 T cells followed by PCR amplification, Sanger sequencing, and decomposition analysis. *B-E*, T cells recovered on day 25 were placed into the Seahorse metabolic analyzer, where the OCR (**B**) was measured simultaneously with ECAR (**C**). Response to oligomycin (**D**) and maximal ECAR values (**E**) were calculated as in Figure 1. Plots in B-E represent data from 2 pooled samples per group (3-4 mice in each pool, 6-8 mice total) and each experiment was performed at least twice. ***p<0.001.

### Decreased glycolytic enzyme activity in AMPK KO T cells

To understand a mechanistic role for AMPK in T cell glycolysis, we set out to identify protein targets of AMPK phosphorylation in GVHD T cells. To accomplish this identification, we utilized an antibody which detects phosphorylation of the AMPK-specific motif LxRxx(pS/pT) to specifically immunoprecipitate proteins phosphorylated by AMPK from day 7 murine T cells. Bulk proteins immunoprecipitated with the phospho-AMPK motif specific antibody were then analyzed by LC-MS to discriminate targets represented at 3-fold or greater levels in WT versus AMPK KO donor cells. This method identified 134 proteins differentially recovered from WT versus AMPK KO T cells on day 7 post-transplant (**Figure 5A, Supp. Table 1**). Consistent with our metabolic results, four of the top candidate proteins represented enzymes involved in glycolytic metabolism, including aldolase A, glyceraldehyde-3-phosphate dehydrogenase (GAPDH), pyruvate kinase M1 (PKM), and enolase (**Figure 5B**). Importantly, there was no difference in total protein levels of these differentially recovered enzymes on day 7 post-transplant prior to immunoprecipitation (**Figure 5C**). Previous literature has demonstrated a role for post-translation modifications in controlling metabolic enzyme activity. To evaluate whether AMPK phosphorylation impacted enzyme function, we quantitate the specific activity of the two most differentially recovered candidates, aldolase and GAPDH. This quantitation was particularly relevant for aldolase, as it was the only precipitated glycolytic enzyme that contained a canonical LxRxx(pS/pT) AMPK phosphorylation motif, potentially identifying it as a direct target of AMPK. Enzyme analysis showed a marked reduction in aldolase activity in AMPK KO T cells following *in vitro* anti-CD3/CD28 stimulation (**Figure 5D-E**). Further, while GAPDH activity remained unchanged following *in vitro* activation (dns), GAPDH activity consistently decreased >20% in AMPK KO T cells on day 7 post-transplant (**Figure 5F**). Together, these data demonstrate that despite equivalent total protein levels, multiple glycolytic enzymes were immunoprecipitated to a lesser degree with an AMPK phospho-motif specific antibody in AMPK KO compared to WT T cells, coincident with decreased enzyme function in two of the most differentially recovered candidates.

**Figure 5.**
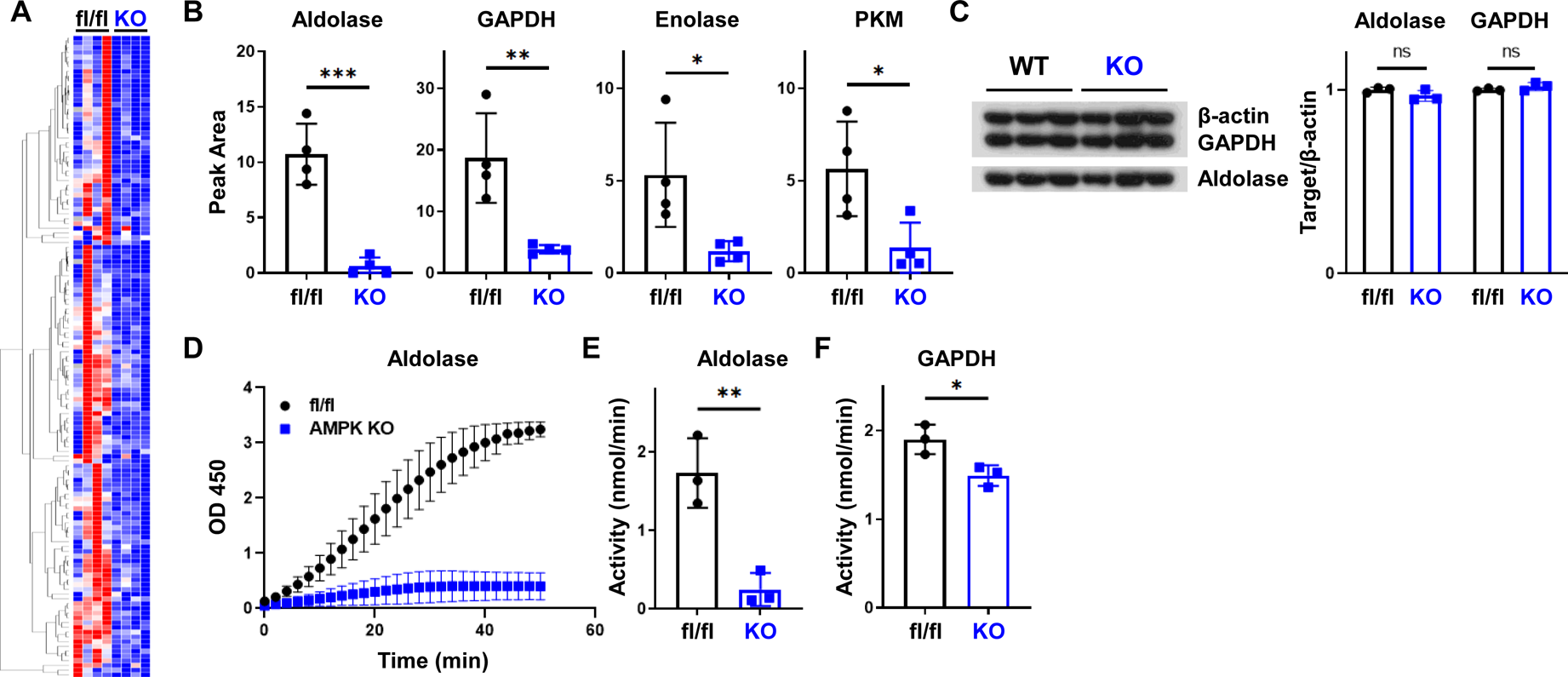
Decreased aldolase activity in AMPK KO T cells. WT versus AMPK KO T cells were transplanted into B6D2F1 recipients, recovered on day 7, and proteins immunoprecipitated from cell lysates using an antibody against the phosphorylated AMPK-specific motif LxRxx(pS/pT). Precipitated candidate proteins were subsequently identified via LC-MS and a heatmap generated of those recovered at 3-fold or higher levels in WT versus AMPK KO T cells (**A**). Representative LC-MS data from the heatmap in *A* is shown for the four glycolytic enzymes of interest (**B**). Control protein samples were recovered from day 7 samples prior to immunoprecipitation and blotted for total levels of candidate proteins aldolase and GAPDH (**C**). *D-E*, WT or AMPK KO T cells were stimulated on CD3/CD28 coated plates for 72 hours, followed by T cell recovery and measurement of aldolase activity in cell lysates. GAPDH was measured in a similar fashion from T cells recovered on day 7 post-transplant (**F**). For data in A-B, 12x WT and 12x AMPK KO T cells were recovered on day 7 post-transplant and divided into four groups of three recipients each. These four groups were then processed as individual replicates through cell lysis, IP, and LC-MS analysis. In C and F, n=3 replicates pooled from 9 or more individual recipients (e.g. 3 groups of 3 recipients each). Graphs in D-E represent data from three, separate biological donors. *p<0.05, **p<0.01, ***p<0.001.

### Decreased glycolysis correlates with decreased IFN**_γ_** production in AMPK KO T cells

Previous work has demonstrated a mechanistic link between decreased glycolysis and decreased IFNγ levels in activated T cells, where diminished IFNγ production resulted from either decreased levels of acetate or a moonlighting role of GAPDH in sequestering IFNγ transcripts ^25, 26^. Given the importance of IFNγ in GVHD pathogenesis, coupled with the impaired glycolysis of AMPK deficient murine and human T cells, we assessed whether a similar association between GAPDH activation and IFNγ levels was operational in AMPK KO T cells. T cells pre-loaded with CellTrace Violet were recovered from spleens of recipient animals on day 7, re-stimulated with F1 splenocytes in the presence of Brefeldin A, and cytokine production measured 6 hours later via intracellular cytokine staining. Significantly fewer AMPK KO cells expressed IFNγ following this re-stimulation compared to WT cells (**Figure 6A-B**). Furthermore, the few AMPK KO T cells that were IFNγ+ demonstrated a markedly reduced median fluorescence intensity, indicating less IFNγ being produced per cell (**Figure 6C**). This reduction in IFNγ production was also seen in AMPK KO cells *in vitro* following stimulation in a mixed leukocyte reaction (**Supp Figure 1A-C**). These IFNγ results are consistent with studies by Lepez et al., who documented decreased IFNγ production by AMPK KO T cells both *in vitro* and in response to homeostatic proliferation *in vivo*^35^. Importantly, absence of AMPK did not globally reduce proinflammatory cytokine production, as AMPK KO T cells generated equivalent amounts of TNF *in vitro* (**Supp Figure 1D-F**) and day 7 AMPK KO T cells had low but equal levels of both TNF and IL-17 in response to F1 splenocyte re-exposure (**Supp Figure 1G** and dns). To understand why IFNγ levels decreased in AMPK KO T cells post-transplant, we recovered RNA from CD4 and CD8 donor T cells on day 7 and performed quantitative RT-PCR. Tellingly, IFNγ transcript levels in both CD4 and CD8 subpopulations were equivalent between WT and AMPK KO T cells on Day 7 (**Supp Figure 1H**), suggesting the lesion occurs at the level of translation and not transcription ^26^. To support this hypothesis, we set out to determine whether changes in IFNγ were cell intrinsic, as would be expected if the intracellular metabolic machinery were influencing the amount of cytokine produced, rather than being influenced solely by the surrounding inflammatory milieu. WT and AMPK KO T cells, congenically marked with CD90.1/2 or CD90.2, respectively (**Figure 6D**), were combined in a 1:1 ratio and transplanted into irradiated F1 recipients. Donor T cells were recovered on day 7 and assessed for intracellular production of IFNγ. Recipients transplanted with only WT or AMPK KO T cells served as controls (**6D, left panels**), again demonstrating a marked defect in IFNγ production from animals singly transplanted with AMPK KO T cells. WT T cells transplanted 1:1 with AMPK KO T cells demonstrated a slight but statistically significant decrease in IFNγ+ percentages on day 7 post-transplant, perhaps as a result of decreased local production of proinflammatory cytokines by neighboring AMPK KO cells. Importantly, however, the sharp decrease in IFNγ production seen in singly transplanted KO cells was maintained when AMPK KO cells were co-transplanted with WT cells (**Figure 6E**), demonstrating altogether that AMPK KO T cells produce less IFNγ upon antigenic re-stimulation in a cell-intrinsic manner.

**Figure 6.**
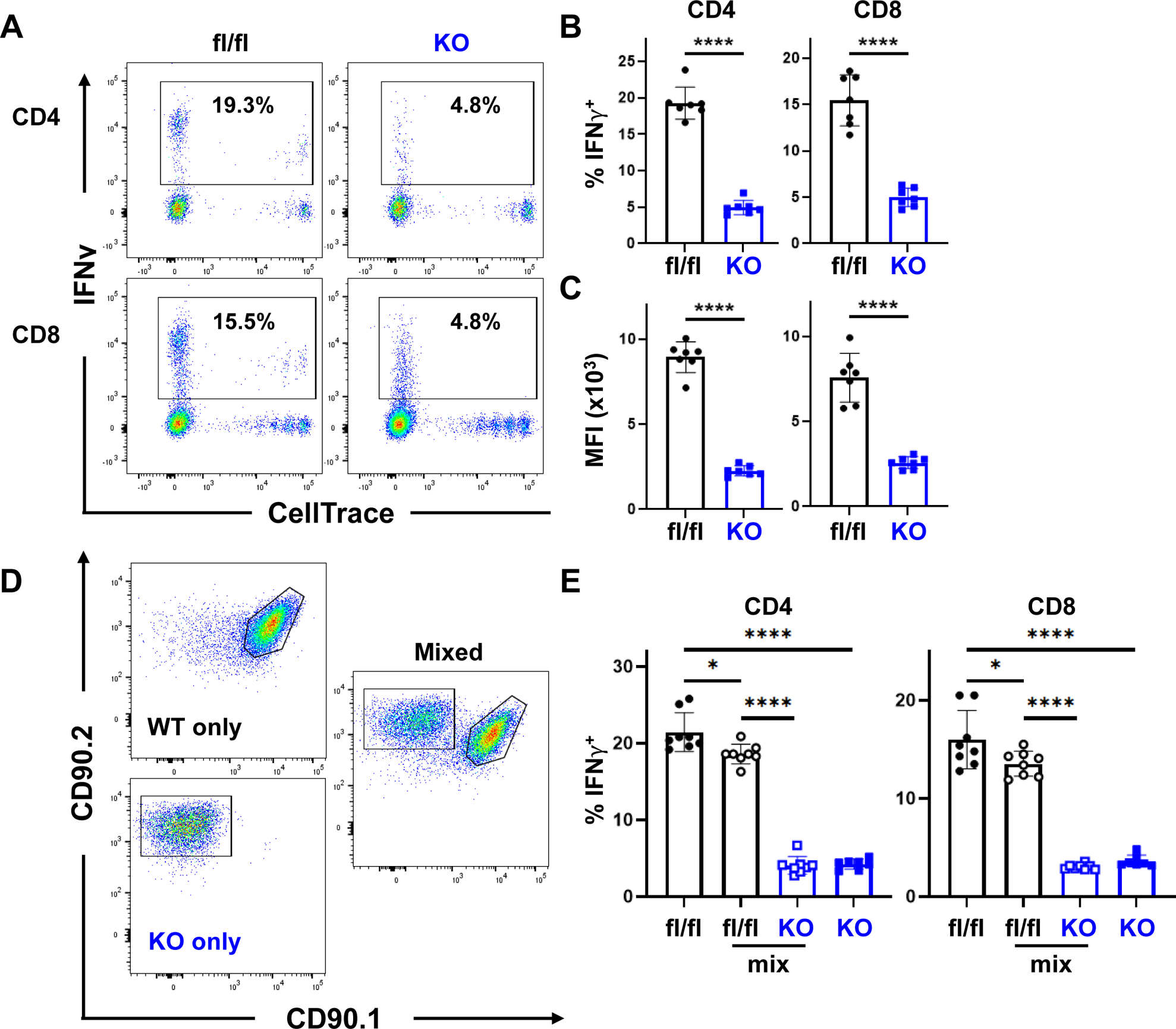
Decreased IFN_γ_ production in AMPK KO T cells is cell-intrinsic. *A-C*, WT or AMPK KO T cells were transplanted individually into irradiated B6D2F1 recipients, recovered on day 7, stimulated for 6 hours with fresh B6D2F1 splenocytes in the presence of Brefeldin A, and analyzed for intracellular cytokine production. Representative flow plots are shown in (**A**), with the percentage of IFNγ+ cells (**B**) and the median fluorescence intensity (MFI) of cells falling within the IFNγ+ gate shown in (**C**), respectively. *D-E*, WT (CD90.1/2) and AMPK KO (CD90.2) T cells were transplanted separately or in a 1:1 combination (mixed) into irradiated B6D2F1 recipients and intracellular IFNγ detected as outlined in A-C. Representative flow plots for individual and mixed samples are shown in (**D**), while (**E**) represents the percentage of IFNγ+ cells in CD4 versus CD8 T cells from multiple samples. n=7-8 recipients/group for all experiments. *p<0.05, ****p<0.0001.

## DISCUSSION

Intracellular metabolism is central to multiple aspects of T cell biology, suggesting that it’s control could be key to ameliorating T cell-mediated disease. We have previously shown that the cellular energy sensor AMPK plays a key role in donor T cells during GVHD initiation and pathogenesis. Elimination of AMPK mitigates GVHD while still preserving homeostatic reconstitution and maintaining beneficial GVL responses. Here we extend these previous studies by documenting the specific changes in metabolism that occur in T cells which lack AMPK, uncovering potential mechanisms that make these changes possible, and correlating metabolic alterations with downstream function. To pave the way for potential clinical application of these findings, we modified a xenogeneic model of GVHD which, when combined with elimination of AMPK in primary human T cells, demonstrated that AMPK is responsible for similar metabolic changes in both murine and human T cells.

Numerous xenogeneic models have been developed over the last 30 years^36^. And while some models replicate GVHD with the transfer of isolated T cells, these approaches often require transgenic expression of growth factors or additional signaling molecules^36, 37^. To avoid these confounding variables, we simply transplanted *in vitro* expanded human T cells alone into NSG mice. However, recovery of donor T cells at early time points in this ‘T cell only’ model was not encouraging. And while clinical GVHD may have eventually developed, the low yield of T cells recovered at three to four weeks post-transplant was not conducive to studying early changes in metabolic reprogramming. Knowing that most NSG models rely on transplantation of a full complement of human PBMCs, we reasoned that absence of non-T cells in our *in vitro* expanded inoculum was likely responsible for the reduced T cell recovery and limited GVHD severity. Indeed, simple re-introduction of 1×10^6^ autologous non-T cells at the time of transplant markedly increased disease severity, as measured by both enhanced T cell recovery and elevated serum IFNγ levels on day 25. While multiple studies have highlighted a role for murine Class I and Class II molecules in driving xenogeneic GVHD^33, 38^, or introduced transgenic human Class II molecules^39, 40^, few reports have detailed the necessity of donor cells beyond T cells in the pre-transplant mix^41^. As suggested by others^34^, we suspect that the addition of autologous non-T cells increases GVHD severity via presentation of murine antigens by human antigen presenting cells^42^, with an attendant increase in T cell activation. These results are also consistent with other xenogeneic systems, including the increases in serum IFNγ, as human T cells are known to develop an effector phenotype in these models^43^. From the host side, CCL-2 is well known to be upregulated in GVHD target organs^44^, particularly at early times post-transplant^45^, and CCR2, the chemokine receptor for CCL-2, plays an important role in the activation and migration of CD8+ T cells into the intestine and liver during GVHD propagation^46^.

Regarding metabolism, we have previously shown that alloreactive T cells upregulate both oxidative and glycolytic metabolism at early times post-transplant^11^. Based on its role in promoting oxidative metabolism in other cell types^47, 48^, we predicted that AMPK would similarly control oxidative metabolism in allogeneic T cells. More surprising was that AMPK seemed to control both oxidative and glycolytic metabolism simultaneously. Glucose restriction is a well-known trigger for AMPK activation^49^ and elegant studies have demonstrated that aldolase facilitates activation of lysosomal pools of AMPK in an AMP/ADP-independent manner. In this context, decreases in the glycolytic intermediate fructose-1,6-bisphosphate frees aldolase from its usual role in glycolysis to instead promote association of AMPK and LKB1 on the lysosomal membrane^50^, which in turn activates AMPK. AMPK could then promote glycolysis directly, through phosphorylation of 6-phosphofructo-2-kinase^51, 52^, or indirectly via phosphorylation of Class II histone deacetylases like HDAC5 ^53^, to increase cell surface expression of glucose transporters. Intriguingly, HDAC5 was identified as a potential phosphorylation target of AMPK in day 7 allogeneic T cells (Supplemental Table 1), although defining the exact role of PFKFB3 and HDAC5 during GVHD remains an active area of investigation.

Significant differences existed in the precipitation of glycolytic enzymes from WT versus AMPK KO T cells, without discernible variation in total protein levels. Furthermore, short of aldolase on Ser176, none of the other preferentially precipitated enzymes contained a canonical LxRxx(pS/pT) AMPK phosphorylation motif. One possible explanation for these results is generation of a supramolecular complex of glycolytic enzymes in activated T cells, one whose association is facilitated by an AMPK phosphorylated target. Indeed, this supramolecular organization has previously been proposed in CD3+ T cells, where enolase, PKM, aldolase, GAPDH, and sirtuin 2 all associate to drive glycolytic function^54^. Our data argue for a similar organization in allogeneic T cells during GVHD. We suspect that AMPK phosphorylates a key target (which could even be Ser176 on aldolase), whose phosphorylation in turn organizes additional glycolytic enzymes into a supramolecular complex, facilitating a more efficient glycolytic response under conditions of oxidative or nutrient stress. Further work is required to fully confirm this hypothesis in activated T cells. Short of this confirmation, it is clear that aldolase activity is sharply reduced in T cells lacking AMPK. To our knowledge, this is the first direct evidence of AMPK controlling aldolase activity. Furthermore, if true, this finding suggests a mutual feedback loop between aldolase, which promotes AMPK activity in the absence of glycolysis^55^, and AMPK, which in turn redirects aldolase activity back towards glycolysis. Additional mechanisms linking AMPK to increased glycolysis could also include AMPK’s phosphorylation of Sirt2, with a subsequent increase in glucose uptake^56^.

AMPK is known to promote IFNγ production in T cells both *in vitro* and in response to homeostatic proliferation *in vivo*^35^. Similar to those studies, we found a muted defect in IFNγ production upon restimulation of day 7 allogeneic T cells, without changes in other proinflammatory cytokines. Disruptions in glycolysis have been previously been linked to diminished IFNγ production, with insufficient acetate production undermining transcription of IFNγ mRNA^25^ and effective IFNγ translation being encumbered by the moonlighting role of GAPDH, which sequesters IFNγ transcripts when glycolysis activity wanes^26^. Our data support a role for disrupted translation as a possible mechanism underlying the decreased IFNγ levels in AMPK KO T cells, a prediction based upon equivalent IFNγ transcript levels in day 7 T cells, reduced immunoprecipitation of GAPDH alongside a number of other glycolytic enzymes, and decreased GAPDH activity levels in day 7 AMPK KO T cells, despite equivalent levels of total GAPDH protein.

T cell-associated IFNγ plays a complex role post-transplant, having previously been ascribed both beneficial and deleterious effects^57–59^. Seminal work by Burman et al. demonstrated an early dependence on IFNγ for preventing overwhelming lung pathology early post-transplant, while T cell-derived IFNγ played a destructive role at later times by promoting a Th1 effector phenotype and increasing inflammation in the gastrointestinal tract ^60^. Our data imply that the production of IFNγ by AMPK KO T cells is sufficient to avoid early lung pathology but may be impactful in decreasing overall GVHD severity when T cells migrate to target tissues. Reassuringly, in spite of decreased IFNγ production, AMPK KO T cells continue to elicit sufficient and potent GVL effects ^14^, a phenotype consistent with recent studies demonstrating that IFNγ production by chimeric antigen receptor T cells *in vivo* is dispensable for controlling leukemia growth^61, 62^.

While the landscape of therapies to successfully prevent GVHD has improved in recent years ^63–66^, further novel strategies are required to maintain or enhance anti-tumor responses while still minimizing GVHD pathogenesis. Human T cells lacking AMPK displayed metabolic vulnerabilities *in vitro* identical to those seen in day 7 murine T cells. However, when administered *in vivo,* AMPK-deficient human T cells continued to manifest impaired glycolysis 25 days later but lost the observed defect in oxidative capacity. This metabolic correction was not due to outgrowth of AMPK positive T cells, as TIDE analysis demonstrated indel rates which continued to exceed 85%. However, multiple alternative explanations might account for these differences between mouse and human T systems: for one, murine cells were analyzed on Day 7 post-transplant, whereas human cells were not collected until Day 25. This change in timing could imply that AMPK has more impact in driving oxidative metabolism in early T cell expansion, with a more sustained influence on glycolytic metabolism over time. Indeed, glycolytic metabolism has been shown to be particularly important in CD8+ T cell memory formation during chronic stimulation^67^, supporting a critical role for maintaining glycolysis at later timepoints in GVHD development. Another possible explanation concerns the duration of AMPK loss. Murine T cells lose AMPK expression during thymic development, then migrate to the periphery and exist for weeks in the absence of AMPK signaling. In contrast, AMPK is acutely deleted in the human model. Sustained loss of AMPK has been previously linked to mitochondrial dysregulation^31^, which means that the murine model may exaggerate the role of AMPK by also impacting long-term mitochondrial capacity. Finally, murine versus human differences could simply reflect the different environmental or experimental conditions between allogeneic and xenogeneic models of GVHD (reviewed in^68^). Regardless, in both murine and human T cells, lesioning AMPK disrupted oxidative metabolism at early timepoints and glycolytic metabolism at all time points, coincident with a reduction in disease severity, highlighting the mechanistic importance of AMPK in GVHD pathogenesis.

In sum, these results demonstrate that AMPK plays a role in both oxidative and glycolytic metabolism during GVHD in both murine and human T cells. Further, our data suggest somewhat surprisingly that the more prominent lesion in human T cells post-transplant was in glycolysis and not oxidative phosphorylation, a result supported by the novel observation that a set of glycolysis-related proteins, are immunoprecipitated together at much lower rates in the absence of AMPK. These findings suggest a model in which AMPK phosphorylation provides a nidus for important interactions among glycolytic enzymes such that in the absence of AMPK, these interactions are lessened, and glycolysis is consequently reduced. Important to our understanding of GVHD, rates of T cell glycolysis can be directly linked to IFNγ production, with a reduction in glycolytic activity freeing GAPDH to sequester additional IFNγ transcripts and thereby limit protein translation. *In vivo*, T cells lacking AMPK simultaneously decrease GAPDH activity and harbor a significant drop in IFNγ production. Together these results document a prominent role for AMPK in controlling T cell glycolysis during GVHD development, offer a mechanistic explanation for the decreased disease severity seen following transplantation of AMPK KO T cells, and provide a strong rationale for further development of AMPK inhibition as a clinically translatable strategy for GVHD prevention and treatment.

## ACKNOWLEDGEMENTS

This work was supported by grants to CAB from the National Institute of Health – NHBLI (K08 HL123631, R01 HL144556), the American Society of Hematology (ASH) Scholar award, and the Be the Match Foundation (Amy Strelzer-Manasevit award). These studies were also made possible by an ASH Research Training Award for Fellows to AR and to a St. Baldrick’s Foundation fellowship and Young Investigator Awards from the Hyundai Motor Company and Alex’s Lemonade Stand Foundation to EB. EB was additionally supported on a University of Pittsburgh Cancer Immunology Training Program T32 grant (5T32CA082084) and an NICHD K12 Grant (HD052892). Cellular bioenergetics was performed on the Seahorse Analyzer in collaboration with the Rangos Metabolic Core within the Department of Pediatrics at the University of Pittsburgh, assisted by Core Director Clinton Van’t Land. The contents of this article are solely the responsibility of the authors and do not necessarily represent the official views of the National Institutes of Health.

## AUTHORSHIP CONTRIBUTIONS

AR designed and performed experiments, analyzed data, and drafted and reviewed the manuscript. ELB designed experiments and wrote/reviewed/edited the manuscript. LKS performed experiments, analyzed data, and drafted the manuscript. DAM designed and performed experiments. CW, MJR, and RC performed experiments. WH designed experiments, analyzed data, generated figures, and drafted the manuscript. CAB drafted the manuscript with assistance from AR, ELB, LKS, and WH, designed and performed experiments, analyzed data, generated figures, and revised the final manuscript. Authorship order (including among co-authors) was assigned based on percent contribution to the final manuscript, overall intellectual involvement, and role in responding to reviewers’ inquiries.

## STUDY APPROVAL

All animal studies were approved and carried out according to Institutional Animal Care and Use Committee guidelines from the University of Pittsburgh. All studies on human cells were designated Exempt status by the University of Pittsburgh Institutional Review Board.

## REFERENCES

1. Zeiser R, Blazar BR. Acute Graft-versus-Host Disease - Biologic Process, Prevention, and Therapy. N. Engl. J. Med. 2017;377(22):2167–2179.

2. Zeiser R, von Bubnoff N, Butler J, et al. Ruxolitinib for Glucocorticoid-Refractory Acute Graft-versus-Host Disease. N. Engl. J. Med. 2020;382(19):1800–1810.

3. Abedin S, McKenna E, Chhabra S, et al. Efficacy, Toxicity, and Infectious Complications in Ruxolitinib-Treated Patients with Corticosteroid-Refractory Graft-versus-Host Disease after Hematopoietic Cell Transplantation. Biol. Blood Marrow Transplant. 2019;25(8):1689–1694.

4. Naesens M, Kuypers DRJ, Sarwal M. Calcineurin inhibitor nephrotoxicity. Clin. J. Am. Soc. Nephrol. 2009;4(2):481–508.

5. Zeiser R, Socié G. The development of ruxolitinib for glucocorticoid-refractory acute graft-versus-host disease. Blood Adv. 2020;4(15):3789–3794.

6. MacMillan ML, Weisdorf DJ, Wagner JE, et al. Response of 443 patients to steroids as primary therapy for acute graft-versus-host disease: comparison of grading systems. Biol. Blood Marrow Transplant. 2002;8(7):387–394.

7. Li J-M, Giver CR, Lu Y, et al. Separating graft-versus-leukemia from graft-versus-host disease in allogeneic hematopoietic stem cell transplantation. Immunotherapy. 2009;1(4):599–621.

8. Rangel Rivera GO, Knochelmann HM, Dwyer CJ, et al. Fundamentals of T cell metabolism and strategies to enhance cancer immunotherapy. Front. Immunol. 2021;12:645242.

9. Reina-Campos M, Scharping NE, Goldrath AW. CD8+ T cell metabolism in infection and cancer. Nat. Rev. Immunol. 2021;21(11):718–738.

10. Elia I, Rowe JH, Johnson S, et al. Tumor cells dictate anti-tumor immune responses by altering pyruvate utilization and succinate signaling in CD8+ T cells. Cell Metab. 2022;34(8):1137–1150.e6.

11. Brown RA, Byersdorfer CA. Metabolic pathways in alloreactive T cells. Front. Immunol. 2020;11:1517.

12. Glick GD, Rossignol R, Lyssiotis CA, et al. Anaplerotic metabolism of alloreactive T cells provides a metabolic approach to treat graft-versus-host disease. J. Pharmacol. Exp. Ther. 2014;351(2):298–307.

13. Gatza E, Wahl DR, Opipari AW, et al. Manipulating the bioenergetics of alloreactive T cells causes their selective apoptosis and arrests graft-versus-host disease. Sci. Transl. Med. 2011;3(67):67ra8.

14. Monlish DA, Beezhold KJ, Chiaranunt P, et al. Deletion of AMPK minimizes graft-versus-host disease through an early impact on effector donor T cells. JCI Insight. 2021;6(14):

15. Nguyen HD, Chatterjee S, Haarberg KMK, et al. Metabolic reprogramming of alloantigen-activated T cells after hematopoietic cell transplantation. J. Clin. Invest. 2016;126(4):1337–1352.

16. Huang Y, Zou Y, Jiao Y, et al. Targeting Glycolysis in Alloreactive T Cells to Prevent Acute Graft-Versus-Host Disease While Preserving Graft-Versus-Leukemia Effect. Front. Immunol. 2022;13:751296.

17. Beezhold K, Moore N, Chiaranunt P, Brown R, Byersdorfer CA. Deletion of AMP-Activated Protein Kinase (AMPK) in Donor T Cells Protects Against Graft-Verus-Host Disease through Control of Regulatory T Cell Expansion and Target Organ Infiltration. Blood. 2016;128(22):806–806.

18. Tamás P, Hawley SA, Clarke RG, et al. Regulation of the energy sensor AMP-activated protein kinase by antigen receptor and Ca2+ in T lymphocytes. J. Exp. Med. 2006;203(7):1665–1670.

19. He N, Fan W, Henriquez B, et al. Metabolic control of regulatory T cell (Treg) survival and function by Lkb1. Proc. Natl. Acad. Sci. USA. 2017;114(47):12542–12547.

20. Willows R, Navaratnam N, Lima A, Read J, Carling D. Effect of different γ-subunit isoforms on the regulation of AMPK. Biochem. J. 2017;474(10):1741–1754.

21. Rabinovitch RC, Samborska B, Faubert B, et al. AMPK Maintains Cellular Metabolic Homeostasis through Regulation of Mitochondrial Reactive Oxygen Species. Cell Rep. 2017;21(1):1–9.

22. Ren Y, Shen H-M. Critical role of AMPK in redox regulation under glucose starvation. Redox Biol. 2019;25:101154.

23. Kazyken D, Magnuson B, Bodur C, et al. AMPK directly activates mTORC2 to promote cell survival during acute energetic stress. Sci. Signal. 2019;12(585):

24. Wang H, Yang Y-G. The complex and central role of interferon-γ in graft-versus-host disease and graft-versus-tumor activity. Immunol. Rev. 2014;258(1):30–44.

25. Peng M, Yin N, Chhangawala S, et al. Aerobic glycolysis promotes T helper 1 cell differentiation through an epigenetic mechanism. Science. 2016;354(6311):481–484.

26. Chang C-H, Curtis JD, Maggi LB, et al. Posttranscriptional control of T cell effector function by aerobic glycolysis. Cell. 2013;153(6):1239–1251.

27. Nakada D, Saunders TL, Morrison SJ. Lkb1 regulates cell cycle and energy metabolism in haematopoietic stem cells. Nature. 2010;468(7324):653–658.

28. Brinkman EK, van Steensel B. Rapid quantitative evaluation of CRISPR genome editing by TIDE and TIDER. Methods Mol. Biol. 2019;1961:29–44.

29. Magee JA, Ikenoue T, Nakada D, et al. Temporal changes in PTEN and mTORC2 regulation of hematopoietic stem cell self-renewal and leukemia suppression. Cell Stem Cell. 2012;11(3):415–428.

30. Wiśniewski JR, Zougman A, Mann M. Combination of FASP and StageTip-based fractionation allows in-depth analysis of the hippocampal membrane proteome. J. Proteome Res. 2009;8(12):5674–5678.

31. Herzig S, Shaw RJ. AMPK: guardian of metabolism and mitochondrial homeostasis. Nat. Rev. Mol. Cell Biol. 2018;19(2):121–135.

32. Ma EH, Poffenberger MC, Wong AH-T, Jones RG. The role of AMPK in T cell metabolism and function. Curr. Opin. Immunol. 2017;46:45–52.

33. King MA, Covassin L, Brehm MA, et al. Human peripheral blood leucocyte non-obese diabetic-severe combined immunodeficiency interleukin-2 receptor gamma chain gene mouse model of xenogeneic graft-versus-host-like disease and the role of host major histocompatibility complex. Clin. Exp. Immunol. 2009;157(1):104–118.

34. Schroeder MA, DiPersio JF. Mouse models of graft-versus-host disease: advances and limitations. Dis. Model. Mech. 2011;4(3):318–333.

35. Lepez A, Pirnay T, Denanglaire S, et al. Long-term T cell fitness and proliferation is driven by AMPK-dependent regulation of reactive oxygen species. Sci. Rep. 2020;10(1):21673.

36. Hess NJ, Brown ME, Capitini CM. GVHD pathogenesis, prevention and treatment: lessons from humanized mouse transplant models. Front. Immunol. 2021;12:723544.

37. Hess NJ, Hudson AW, Hematti P, Gumperz JE. Early T Cell Activation Metrics Predict Graft-versus-Host Disease in a Humanized Mouse Model of Hematopoietic Stem Cell Transplantation. J. Immunol. 2020;205(1):272–281.

38. Brehm MA, Kenney LL, Wiles MV, et al. Lack of acute xenogeneic graft-versus-host disease, but retention of T-cell function following engraftment of human peripheral blood mononuclear cells in NSG mice deficient in MHC class I and II expression. FASEB J. 2019;33(3):3137–3151.

39. Ehx G, Somja J, Warnatz H-J, et al. Xenogeneic Graft-Versus-Host Disease in Humanized NSG and NSG-HLA-A2/HHD Mice. Front. Immunol. 2018;9:1943.

40. Covassin L, Laning J, Abdi R, et al. Human peripheral blood CD4 T cell-engrafted non-obese diabetic-scid IL2rγ(null) H2-Ab1 (tm1Gru) Tg (human leucocyte antigen D-related 4) mice: a mouse model of human allogeneic graft-versus-host disease. Clin. Exp. Immunol. 2011;166(2):269–280.

41. Kawasaki Y, Sato K, Hayakawa H, et al. Comprehensive Analysis of the Activation and Proliferation Kinetics and Effector Functions of Human Lymphocytes, and Antigen Presentation Capacity of Antigen-Presenting Cells in Xenogeneic Graft-Versus-Host Disease. Biol. Blood Marrow Transplant. 2018;24(8):1563–1574.

42. Lucas PJ, Shearer GM, Neudorf S, Gress RE. The human antimurine xenogeneic cytotoxic response. I. Dependence on responder antigen-presenting cells. J. Immunol. 1990;144(12):4548–4554.

43. Ali N, Flutter B, Sanchez Rodriguez R, et al. Xenogeneic graft-versus-host-disease in NOD-scid IL-2Rγnull mice display a T-effector memory phenotype. PLoS One. 2012;7(8):e44219.

44. Mapara MY, Leng C, Kim Y-M, et al. Expression of chemokines in GVHD target organs is influenced by conditioning and genetic factors and amplified by GVHR. Biol. Blood Marrow Transplant. 2006;12(6):623– 634.

45. Castor MGM, Pinho V, Teixeira MM. The role of chemokines in mediating graft versus host disease: opportunities for novel therapeutics. Front. Pharmacol. 2012;3:23.

46. Terwey TH, Kim TD, Kochman AA, et al. CCR2 is required for CD8-induced graft-versus-host disease. Blood. 2005;106(9):3322–3330.

47. Blagih J, Coulombe F, Vincent EE, et al. The energy sensor AMPK regulates T cell metabolic adaptation and effector responses in vivo. Immunity. 2015;42(1):41–54.

48. Yang K, Chi H. AMPK helps T cells survive nutrient starvation. Immunity. 2015;42(1):4–6.

49. Salt IP, Johnson G, Ashcroft SJ, Hardie DG. AMP-activated protein kinase is activated by low glucose in cell lines derived from pancreatic beta cells, and may regulate insulin release. Biochem. J. 1998;335 (Pt 3)(Pt 3):533–539.

50. Zhang C-S, Hawley SA, Zong Y, et al. Fructose-1,6-bisphosphate and aldolase mediate glucose sensing by AMPK. Nature. 2017;548(7665):112–116.

51. Bando H, Atsumi T, Nishio T, et al. Phosphorylation of the 6-phosphofructo-2-kinase/fructose 2,6-bisphosphatase/PFKFB3 family of glycolytic regulators in human cancer. Clin. Cancer Res. 2005;11(16):5784–5792.

52. Doménech E, Maestre C, Esteban-Martínez L, et al. AMPK and PFKFB3 mediate glycolysis and survival in response to mitophagy during mitotic arrest. Nat. Cell Biol. 2015;17(10):1304–1316.

53. McGee SL, van Denderen BJW, Howlett KF, et al. AMP-activated protein kinase regulates GLUT4 transcription by phosphorylating histone deacetylase 5. Diabetes. 2008;57(4):860–867.

54. Hamaidi I, Zhang L, Kim N, et al. Sirt2 Inhibition Enhances Metabolic Fitness and Effector Functions of Tumor-Reactive T Cells. Cell Metab. 2020;32(3):420–436.e12.

55. Lin S-C, Hardie DG. AMPK: Sensing Glucose as well as Cellular Energy Status. Cell Metab. 2018;27(2):299–313.

56. Liemburg-Apers DC, Wagenaars JAL, Smeitink JAM, Willems PHGM, Koopman WJH. Acute stimulation of glucose influx upon mitoenergetic dysfunction requires LKB1, AMPK, Sirt2 and mTOR-RAPTOR. J. Cell Sci. 2016;129(23):4411–4423.

57. Wang H, Asavaroengchai W, Yeap BY, et al. Paradoxical effects of IFN-gamma in graft-versus-host disease reflect promotion of lymphohematopoietic graft-versus-host reactions and inhibition of epithelial tissue injury. Blood. 2009;113(15):3612–3619.

58. Lu Y, Waller EK. Dichotomous role of interferon-gamma in allogeneic bone marrow transplant. Biol. Blood Marrow Transplant. 2009;15(11):1347–1353.

59. Murphy WJ, Welniak LA, Taub DD, et al. Differential effects of the absence of interferon-gamma and IL-4 in acute graft-versus-host disease after allogeneic bone marrow transplantation in mice. J. Clin. Invest. 1998;102(9):1742–1748.

60. Burman AC, Banovic T, Kuns RD, et al. IFNgamma differentially controls the development of idiopathic pneumonia syndrome and GVHD of the gastrointestinal tract. Blood. 2007;110(3):1064–1072.

61. Bailey SR, Vatsa S, Larson RC, et al. Blockade or Deletion of IFNγ Reduces Macrophage Activation without Compromising CAR T-cell Function in Hematologic Malignancies. Blood Cancer Discov. 2022;3(2):136–153.

62. Larson RC, Kann MC, Bailey SR, et al. CAR T cell killing requires the IFNγR pathway in solid but not liquid tumours. Nature. 2022;604(7906):563–570.

63. Watkins B, Qayed M, McCracken C, et al. Phase II trial of costimulation blockade with abatacept for prevention of acute GVHD. J. Clin. Oncol. 2021;39(17):1865–1877.

64. Stenger EO, Watkins B, Rogowski K, et al. Abatacept GVHD prophylaxis in unrelated hematopoietic cell transplantation for pediatric bone marrow failure. Blood Adv. 2023;

65. O’Donnell PV, Luznik L, Jones RJ, et al. Nonmyeloablative bone marrow transplantation from partially HLA-mismatched related donors using posttransplantation cyclophosphamide. Biol. Blood Marrow Transplant. 2002;8(7):377–386.

66. Al-Homsi AS, Roy TS, Cole K, Feng Y, Duffner U. Post-transplant high-dose cyclophosphamide for the prevention of graft-versus-host disease. Biol. Blood Marrow Transplant. 2015;21(4):604–611.

67. Phan AT, Doedens AL, Palazon A, et al. Constitutive Glycolytic Metabolism Supports CD8+ T Cell Effector Memory Differentiation during Viral Infection. Immunity. 2016;45(5):1024–1037.

68. Koyama M, Hill GR. Mouse models of antigen presentation in hematopoietic stem cell transplantation. Front. Immunol. 2021;12:715893.

69. Brinkman EK, Chen T, Amendola M, van Steensel B. Easy quantitative assessment of genome editing by sequence trace decomposition. Nucleic Acids Res. 2014;42(22):e168.

