## Supplemental pdf for "AMPK Drives Both Glycolytic and Oxidative Metabolism in T Cells During Graft-versus-host Disease"

### Supplemental Materials

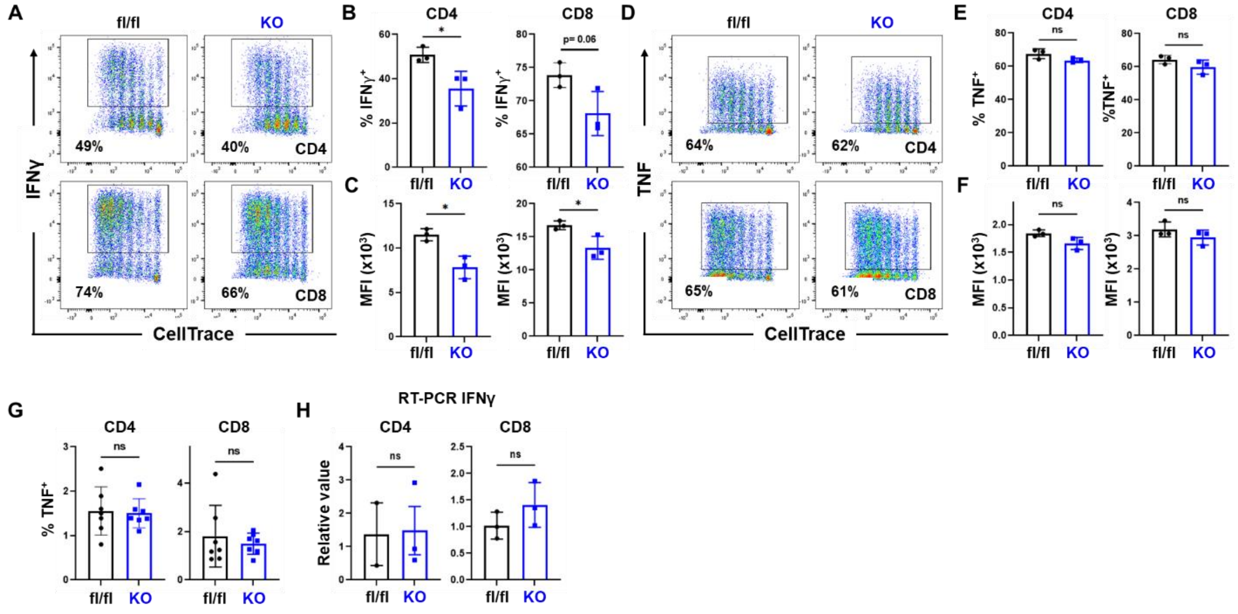

**Supplemental Figure 1. Decreased IFNγ production in AMPK KO T cells stimulated *in vitro*** *A-C*, WT versus AMPK KO were labeled with CellTrace Violet and placed into a mixed leukocyte reaction with B6D2F1 splenocytes. At 72 hours, cells were recovered and re-stimulated on anti-CD3/CD28 antibody coated plates for 4 hours in the presence of Brefeldin A, followed by analysis of intracellular cytokine production. Representative flow plots are shown in (**A**), with the percentage of IFNγ+ cells (**B**) and the median fluorescence intensity (MFI) of IFNγ+ in positive cells shown in (**C**), respectively. A similar analysis was done for TNF production (**D-F**). *G*, WT versus AMPK KO were transplanted into irradiated B6D2F1 recipients, recovered on day 7, and stimulated for 6 hours with fresh B6D2F1 splenocytes in the presence of Brefeldin A, followed by analysis of intracellular TNF. The percentage of TNF+ cells for both CD4 and CD8 T cells is shown. *H*, WT versus AMPK KO were labeled with CellTrace Violet and transplanted into irradiated B6D2F1 recipients. On day 7 post-transplant, CD4 and CD8 T donor T cells undergoing more than 8 divisions were recovered and RNA extracted. Quantitative RT-PCR then measured levels of IFNγ transcript relative to values present in WT cells. A-F, n= 3 biological replicates per group. In G, n=7-8 replicates/group. For H, n=3 groups pooled from a total of 9-10 recipients (e.g. 3 groups of 3 recipients each). *$p<0.05$

| Accession | NAME | Significance | wt1 Area | wt2 Area | wt3 Area | wt4 Area | ko1 Area | ko2 Area | ko3 Area | ko4 Area | Description |
|---|---|---|---|---|---|---|---|---|---|---|---|
| P07356 | ANXA2 | 19.06 | 1.90E+05 | 1.25E+06 | 2.72E+05 | 5.73E+05 | 9.96E+04 | 1.81E+05 | 4.94E+04 | 1.67E+05 | Annexin A2 OS=Mus musculus GN=Anxa2 PE=1 SV=2 |
| Q32M21 | GSDA2 | 18.82 | 2.86E+03 | 5.63E+04 | 1.59E+03 | 1.68E+05 | 1000 | 3.81E+03 | 1000 | 1000 | Gasdermin-A2 OS=Mus musculus GN=Gsdma2 PE=2 SV=1 |
| P20029 | GRP78 | 18.41 | 1.29E+04 | 9.33E+04 | 8.11E+04 | 2.55E+04 | 1000 | 6.41E+03 | 1.07E+04 | 5.97E+03 | 78 kDa glucose-regulated protein OS=Mus musculus GN=Hspa5 PE=1 SV=3 |
| E9Q557 | DESP | 18.17 | 2.81E+04 | 2.85E+05 | 7.55E+04 | 4.09E+04 | 6.70E+03 | 5.60E+04 | 4.83E+03 | 1.47E+04 | Desmoplakin OS=Mus musculus GN=Dsp PE=1 SV=1 |
| Q9JLF6 | TGM1 | 17.83 | 1000 | 7.59E+04 | 1.06E+04 | 9.51E+04 | 1000 | 2.71E+03 | 1.03E+04 | 1000 | Protein-glutamine gamma-glutamyltransferase K OS=Mus musculus GN=Tgm1 PE=1 SV=2 |
| Q61233 | PLSL | 16.87 | 1.98E+04 | 3.02E+04 | 8.92E+04 | 3.91E+04 | 1.54E+04 | 1.39E+04 | 1.30E+04 | 1000 | Plastin-2 OS=Mus musculus GN=Lcp1 PE=1 SV=4 |
| P17182 | ENOA | 16.04 | 3.18E+04 | 1.54E+05 | 4.92E+04 | 3.75E+04 | 1.70E+04 | 1.57E+04 | 8.29E+03 | 5.95E+03 | Alpha-enolase OS=Mus musculus GN=Eno1 PE=1 SV=3 |
| P17897 | LYZ1 | 13.22 | 6.91E+04 | 5.83E+04 | 1.89E+05 | 5.97E+04 | 1.06E+04 | 1.40E+04 | 9.65E+03 | 1000 | Lysozyme C-1 OS=Mus musculus GN=Lyz1 PE=1 SV=1 |
| Q921M3 | SF3B3 | 13 | 6.68E+04 | 1.58E+05 | 6.40E+04 | 7.52E+04 | 2.94E+04 | 2.18E+04 | 1.95E+04 | 2.79E+04 | Splicing factor 3B subunit 3 OS=Mus musculus GN=Sf3b3 PE=1 SV=1 |
| P35700 | PRDX1 | 12.43 | 6.65E+04 | 1.00E+05 | 7.99E+04 | 5.85E+04 | 2.25E+04 | 2.55E+04 | 1.99E+04 | 1.31E+04 | Peroxiredoxin-1 OS=Mus musculus GN=Prdx1 PE=1 SV=1 |
| P11499 | HS90B | 11.56 | 7.08E+03 | 1.07E+05 | 1.33E+05 | 3.89E+04 | 3.70E+04 | 1.56E+04 | 3.32E+04 | 5.13E+03 | Heat shock protein HSP 90-beta OS=Mus musculus GN=Hsp90ab1 PE=1 SV=3 |
| Q00941 | CSF2R | 11.45 | 7.61E+05 | 8.22E+05 | 1.09E+05 | 2.19E+06 | 1.16E+05 | 3.22E+05 | 1.25E+05 | 1.79E+05 | Granulocyte-macrophage colony-stimulating factor receptor subunit alpha OS=Mus musculus GN=Csf2ra PE=2 SV=2 |
| Q3UEB3 | PUF60 | 11.34 | 6.59E+05 | 8.21E+05 | 3.92E+05 | 1.72E+06 | 6.49E+04 | 5.33E+05 | 7.21E+04 | 1.49E+05 | Poly(U)-binding-splicing factor PUF60 OS=Mus musculus GN=Puf60 PE=1 SV=2 |
| P12970 | RL7A | 10.58 | 5.63E+04 | 1.41E+05 | 1.57E+05 | 4.19E+05 | 7.70E+04 | 6.00E+04 | 5.45E+04 | 3.37E+04 | 60S ribosomal protein L7a OS=Mus musculus GN=Rpl7a PE=1 SV=2 |
| P26041 | MOES | 10.12 | 1000 | 2.03E+04 | 5.39E+04 | 8.95E+04 | 1000 | 1000 | 1.37E+04 | 8.51E+03 | Moesin OS=Mus musculus GN=Msn PE=1 SV=3 |
| Q3THE2 | ML12B | 10.01 | 3.09E+05 | 5.13E+05 | 6.84E+05 | 5.98E+05 | 1.70E+05 | 1.72E+05 | 1.41E+05 | 9.17E+04 | Myosin regulatory light chain 12B OS=Mus musculus GN=Myl12b PE=1 SV=2 |
| Q9QXL1 | KI21B | 9.99 | 8.60E+04 | 1.78E+06 | 6.60E+05 | 2.51E+06 | 1.58E+05 | 6.61E+05 | 3.62E+05 | 2.20E+05 | Kinesin-like protein KIF21B OS=Mus musculus GN=Kif21b PE=1 SV=2 |
| P29341 | PABP1 | 9.74 | 2.45E+04 | 1.21E+05 | 1.23E+05 | 7.19E+04 | 9.72E+03 | 2.76E+04 | 1.79E+04 | 2.97E+04 | Polyadenylate-binding protein 1 OS=Mus musculus GN=Pabpc1 PE=1 SV=2 |
| P10126 | EF1A1 | 9.15 | 3.23E+04 | 2.83E+05 | 1.78E+05 | 1000 | 4.80E+04 | 3.94E+04 | 3.17E+04 | 2.32E+04 | Elongation factor 1-alpha 1 OS=Mus musculus GN=Eef1a1 PE=1 SV=3 |
| P62242 | RS8 | 8.88 | 1.29E+05 | 2.57E+05 | 2.02E+05 | 1.74E+05 | 8.24E+04 | 1.11E+05 | 9.51E+04 | 4.44E+04 | 40S ribosomal protein S8 OS=Mus musculus GN=Rps8 PE=1 SV=2 |
| Q80U16 | FA65B | 8.88 | 1.25E+05 | 1.32E+05 | 1.21E+05 | 1.58E+05 | 5.85E+04 | 6.07E+04 | 7.06E+04 | 4.43E+04 | Protein FAM65B OS=Mus musculus GN=Fam65b PE=1 SV=2 |
| Q8C115 | PKHH2 | 8.86 | 6.86E+04 | 2.93E+06 | 1.18E+05 | 1.58E+05 | 1.73E+04 | 6.27E+04 | 1000 | 4.55E+04 | Pleckstrin homology domain-containing family H member 2 OS=Mus musculus GN=Plekhh2 PE=1 SV=3 |
| Q7TN02 | MED26 | 8.8 | 1000 | 1.01E+06 | 8.10E+04 | 1.81E+06 | 1000 | 2.02E+05 | 1000 | 1000 | Mediator of RNA polymerase II transcription subunit 26 OS=Mus musculus GN=Med26 PE=1 SV=1 |
| P18760 | COF1 | 8.76 | 2.51E+04 | 2.35E+05 | 2.99E+05 | 2.54E+05 | 3.85E+04 | 3.23E+04 | 1.86E+04 | 1.16E+04 | Cofilin-1 OS=Mus musculus GN=Cfl1 PE=1 SV=3 |
| Q9CXF4 | TBC15 | 8.71 | 3.54E+04 | 8.29E+04 | 3.45E+04 | 2.37E+05 | 4.59E+03 | 1.24E+04 | 1.55E+04 | 1.22E+04 | TBC1 domain family member 15 OS=Mus musculus GN=Tbc1d15 PE=1 SV=1 |
| P05064 | ALDOA | 8.69 | 9.37E+04 | 1.11E+05 | 8.00E+04 | 1.44E+05 | 1.69E+04 | 7.08E+03 | 1000 | 1000 | Fructose-bisphosphate aldolase A OS=Mus musculus GN=Aldoa PE=1 SV=2 |
| Q08943 | SSRP1 | 8.64 | - | 1.49E+05 | 2.61E+04 | 1000 | 4.40E+03 | 1000 | 1000 | 1000 | FACT complex subunit SSRP1 OS=Mus musculus GN=Ssrp1 PE=1 SV=2 |
| P62908 | RS3 | 8.58 | 1.60E+05 | 2.68E+05 | 2.33E+05 | 2.44E+05 | 1.00E+05 | 1.13E+05 | 1.28E+05 | 8.17E+04 | 40S ribosomal protein S3 OS=Mus musculus GN=Rps3 PE=1 SV=1 |
| Q9Z0U1 | ZO2 | 8.11 | 3.75E+04 | 7.14E+04 | 7.46E+04 | 1.03E+05 | 2.77E+04 | 2.48E+04 | 2.18E+04 | 4.35E+04 | Tight junction protein ZO-2 OS=Mus musculus GN=Tjp2 PE=1 SV=2 |
| Q3UHD9 | AGAP2 | 8.01 | 3.63E+04 | 3.72E+04 | 5.48E+04 | 9.39E+04 | 1.97E+04 | 2.92E+04 | 2.84E+04 | 1.37E+04 | Arf-GAP with GTPase ANK repeat and PH domain-containing protein 2 OS=Mus musculus GN=Agap2 PE=1 SV=1 |
| Q61210 | ARHG1 | 8.01 | 4.97E+05 | 3.34E+05 | 4.84E+05 | 6.06E+05 | 2.16E+05 | 2.87E+05 | 1.86E+05 | 2.59E+05 | Rho guanine nucleotide exchange factor 1 OS=Mus musculus GN=Arhgef1 PE=1 SV=2 |
| P63268 | ACTH | 7.86 | 1.50E+05 | 3.25E+05 | 2.86E+05 | 2.11E+05 | 9.87E+04 | 6.02E+04 | 1.09E+05 | 7.54E+04 | Actin gamma-enteric smooth muscle OS=Mus musculus GN=Actg2 PE=1 SV=1 |
| Q5U4C3 | SFR19 | 7.82 | 1000 | 6.74E+04 | 7.57E+04 | 1.17E+05 | 1000 | 1.97E+04 | 1000 | 1.79E+04 | Splicing factor arginine/serine-rich 19 OS=Mus musculus GN=Scaf1 PE=1 SV=1 |
| Q5DU56 | NLRC3 | 7.8 | 1.57E+06 | 2.67E+06 | 6.35E+05 | 2.58E+06 | 1.40E+05 | 4.92E+05 | 1.46E+05 | 3.55E+05 | Protein NLRC3 OS=Mus musculus GN=Nlrc3 PE=2 SV=2 |
| Q8VH51 | RBM39 | 7.64 | 1.40E+05 | 1.34E+05 | 1.24E+05 | 1.71E+05 | 7.61E+04 | 8.88E+04 | 6.54E+04 | 4.45E+04 | RNA-binding protein 39 OS=Mus musculus GN=Rbm39 PE=1 SV=2 |
| P57776 | EF1D | 7.59 | - | 1.01E+05 | 7.19E+04 | 1.67E+05 | 1.17E+04 | 3.56E+04 | 5.64E+03 | 1.55E+04 | Elongation factor 1-delta OS=Mus musculus GN=Eef1d PE=1 SV=3 |
| Q61656 | DDX5 | 7.57 | 1.30E+05 | 1.36E+05 | 1.40E+05 | 1.34E+05 | 6.47E+04 | 7.22E+04 | 8.39E+04 | 3.58E+04 | Probable ATP-dependent RNA helicase DDX5 OS=Mus musculus GN=Ddx5 PE=1 SV=2 |
| P14131 | RS16 | 7.39 | 5.16E+04 | 7.26E+04 | 4.93E+04 | 4.70E+04 | 2.27E+04 | 2.81E+04 | 2.41E+04 | 2.86E+04 | 40S ribosomal protein S16 OS=Mus musculus GN=Rps16 PE=1 SV=4 |
| P63017 | HSP7C | 7.36 | 4.53E+04 | 9.02E+04 | 1.97E+05 | 5.00E+04 | 4.76E+04 | 4.65E+04 | 4.28E+04 | 2.25E+04 | Heat shock cognate 71 kDa protein OS=Mus musculus GN=Hspa8 PE=1 SV=1 |
| P16858 | G3P | 7.36 | 1.59E+04 | 1.21E+05 | 2.90E+05 | 7.61E+04 | 3.85E+04 | 4.72E+04 | 3.70E+04 | 3.04E+04 | Glyceraldehyde-3-phosphate dehydrogenase OS=Mus musculus GN=Gapdh PE=1 SV=2 |
| Q61097 | KSR1 | 7.31 | 1000 | 5.56E+04 | 3.79E+04 | 1.35E+05 | 8.65E+03 | 2.75E+04 | 1000 | 1000 | Kinase suppressor of Ras 1 OS=Mus musculus GN=Ksr1 PE=1 SV=1 |
| P62984 | RL40 | 7.2 | 5.62E+05 | 1.25E+06 | 1.05E+06 | 9.58E+05 | 4.52E+05 | 6.84E+05 | 4.31E+05 | 2.82E+05 | Ubiquitin-60S ribosomal protein L40 OS=Mus musculus GN=Uba52 PE=1 SV=2 |
| Q8R3N1 | NOP14 | 7.19 | 6.81E+05 | 1.48E+06 | 4.03E+05 | 1.57E+06 | 7.01E+04 | 4.75E+05 | 1.01E+05 | 4.54E+04 | Nucleolar protein 14 OS=Mus musculus GN=Nop14 PE=1 SV=2 |
| O89053 | COR1A | 7.12 | 2.16E+04 | 3.04E+04 | 9.50E+04 | 2.59E+04 | 2.64E+04 | 2.05E+04 | 1.67E+04 | 7.85E+03 | Coronin-1A OS=Mus musculus GN=Coro1a PE=1 SV=5 |
| Q6ZWQ0 | SYNE2 | 7.02 | 5.80E+05 | 7.06E+05 | 3.80E+05 | 2.59E+06 | 7.81E+04 | 3.99E+05 | 7.06E+04 | 1.91E+05 | Nesprin-2 OS=Mus musculus GN=Syne2 PE=1 SV=2 |
| P62281 | RS11 | 6.92 | 7.15E+04 | 2.09E+05 | 1.15E+05 | 1.19E+05 | 4.49E+04 | 5.28E+04 | 5.44E+04 | 4.20E+04 | 40S ribosomal protein S11 OS=Mus musculus GN=Rps11 PE=1 SV=3 |
| Q6ZQK5 | ACAP2 | 6.8 | 7.43E+04 | 2.90E+05 | 7.97E+04 | 1.47E+05 | 1.39E+04 | 4.45E+04 | 1.11E+04 | 3.47E+04 | Arf-GAP with coiled-coil ANK repeat and PH domain-containing protein 2 OS=Mus musculus GN=Acap2 PE=1 SV=2 |
| P62245 | RS15A | 6.78 | 8.88E+04 | 1.03E+05 | 1.96E+05 | 1.08E+05 | 2.50E+04 | 6.55E+04 | 3.18E+04 | 3.32E+04 | 40S ribosomal protein S15a OS=Mus musculus GN=Rps15a PE=1 SV=2 |
| P97351 | RS3A | 6.73 | 2.27E+04 | 1.08E+05 | 9.33E+04 | 7.98E+04 | 3.75E+04 | 4.14E+04 | 4.48E+04 | 1.07E+04 | 40S ribosomal protein S3a OS=Mus musculus GN=Rps3a PE=1 SV=3 |
| Q99NE5 | RIMS1 | 6.56 | 1.03E+05 | 4.06E+05 | 1.16E+05 | 2.89E+05 | 2.32E+04 | 8.39E+04 | 1.70E+04 | 4.49E+04 | Regulating synaptic membrane exocytosis protein 1 OS=Mus musculus GN=Rims1 PE=1 SV=2 |
| Q6QI06 | RICTR | 6.56 | 7.43E+05 | 9.22E+05 | 3.82E+05 | 1.22E+06 | 7.55E+04 | 2.33E+05 | 4.85E+04 | 1.14E+05 | Rapamycin-insensitive companion of mTOR OS=Mus musculus GN=Rictor PE=1 SV=2 |
| Q61315 | APC | 6.54 | 1000 | 1.28E+05 | 6.23E+04 | 7.93E+04 | 1000 | 3.89E+04 | 1000 | 1.03E+04 | Adenomatous polyposis coli protein OS=Mus musculus GN=Apc PE=1 SV=1 |
| Q9JJW6 | ALRF2 | 6.42 | 7.11E+03 | 3.32E+04 | 7.56E+04 | 4.17E+04 | 1.27E+04 | 4.06E+04 | 1.26E+04 | 5.42E+03 | Aly/REF export factor 2 OS=Mus musculus GN=Alyref2 PE=1 SV=1 |
| Q8R502 | LRC8C | 6.34 | 1.52E+05 | 2.48E+05 | 1.58E+05 | 1.25E+05 | 8.08E+04 | 9.44E+04 | 8.89E+04 | 7.23E+04 | Volume-regulated anion channel subunit LRRC8C OS=Mus musculus GN=Lrrc8c PE=1 SV=1 |
| P14115 | RL27A | 6.29 | 2.41E+05 | 1.09E+05 | 1.55E+05 | 1.48E+05 | 5.62E+04 | 9.30E+04 | 7.20E+04 | 3.88E+04 | 60S ribosomal protein L27a OS=Mus musculus GN=Rpl27a PE=1 SV=5 |
| P84104 | SRSF3 | 6.27 | 6.99E+04 | 1.14E+05 | 8.75E+04 | 1.02E+05 | 3.09E+04 | 3.64E+04 | 4.70E+04 | 3.11E+04 | Serine/arginine-rich splicing factor 3 OS=Mus musculus GN=Srsf3 PE=1 SV=1 |
| P20152 | VIME | 6.25 | 2.94E+05 | 4.58E+05 | 3.11E+05 | 1.98E+05 | 8.08E+04 | 1.03E+05 | 2.09E+05 | 1.07E+05 | Vimentin OS=Mus musculus GN=Vim PE=1 SV=3 |
| P59999 | ARPC4 | 6.25 | 4.72E+04 | 4.78E+04 | 7.81E+04 | 4.97E+04 | 1.15E+04 | 1.20E+04 | 6.77E+03 | 1.27E+04 | Actin-related protein 2/3 complex subunit 4 OS=Mus musculus GN=Arpc4 PE=1 SV=3 |
| P62858 | RS28 | 6.22 | 2.89E+04 | 5.47E+04 | 6.55E+04 | 1.47E+05 | 1.19E+04 | 2.18E+04 | 1.59E+04 | 7.97E+03 | 40S ribosomal protein S28 OS=Mus musculus GN=Rps28 PE=1 SV=1 |
| Q9Z2V6 | HDAC5 | 6.21 | 2.16E+04 | 4.49E+05 | 1.56E+05 | 4.69E+04 | 4.35E+04 | 5.46E+04 | 2.37E+04 | 1.06E+04 | Histone deacetylase 5 OS=Mus musculus GN=Hdac5 PE=1 SV=2 |
| Q3TBD2 | HMHA1 | 6.15 | 1.78E+04 | 2.01E+04 | 6.47E+04 | 5.01E+04 | 1.67E+04 | 3.03E+04 | 1.18E+04 | 1.37E+04 | Minor histocompatibility protein HA-1 OS=Mus musculus GN=Hmha1 PE=1 SV=2 |
| P51881 | ADT2 | 6.15 | 4.44E+04 | 5.45E+04 | 7.87E+04 | 7.53E+04 | 3.00E+04 | 5.07E+04 | 2.30E+04 | 1.73E+04 | ADP/ATP translocase 2 OS=Mus musculus GN=Slc25a5 PE=1 SV=3 |
| P62918 | RL8 | 6.12 | - | 1.39E+05 | 1.33E+05 | 1.16E+05 | 4.43E+04 | 9.23E+04 | 4.25E+04 | 2.90E+04 | 60S ribosomal protein L8 OS=Mus musculus GN=Rpl8 PE=1 SV=2 |
| P21107 | TPM3 | 6.03 | 6.44E+04 | 1.38E+05 | 1.74E+05 | 8.70E+04 | 8.02E+04 | 5.01E+04 | 4.05E+04 | 2.11E+04 | Tropomyosin alpha-3 chain OS=Mus musculus GN=Tpm3 PE=1 SV=3 |
| Q3U108 | ARI5A | 6.02 | 1.49E+06 | 7.34E+06 | 2.11E+06 | 1.26E+07 | 4.92E+05 | 2.91E+06 | 5.43E+05 | 9.98E+05 | AT-rich interactive domain-containing protein 5A OS=Mus musculus GN=Arid5a PE=1 SV=1 |
| P62204 | CALM | 5.95 | 5.84E+04 | 1.13E+05 | 1.33E+05 | 6.38E+04 | 4.72E+04 | 4.35E+04 | 3.41E+04 | 2.37E+04 | Calmodulin OS=Mus musculus GN=Calm1 PE=1 SV=2 |
| Q9R1C7 | PR40A | 5.91 | 6.32E+04 | 8.03E+04 | 7.95E+04 | 8.10E+04 | 3.14E+04 | 3.25E+04 | 3.29E+04 | 2.93E+04 | Pre-mRNA-processing factor 40 homolog A OS=Mus musculus GN=Prpf40a PE=1 SV=1 |
| P62960 | YBOX1 | 5.65 | 4.65E+04 | 6.42E+04 | 8.81E+04 | 4.74E+04 | 2.89E+04 | 3.47E+04 | 3.51E+04 | 1.73E+04 | Nuclease-sensitive element-binding protein 1 OS=Mus musculus GN=Ybx1 PE=1 SV=3 |
| Q80TI1 | PKHH1 | 5.61 | 3.06E+05 | 1.22E+06 | 4.39E+05 | 2.25E+06 | 1.25E+05 | 5.90E+05 | 7.99E+04 | 1.56E+05 | Pleckstrin homology domain-containing family H member 1 OS=Mus musculus GN=Plekhh1 PE=1 SV=2 |
| P53026 | RL10A | 5.43 | 1000 | 1.06E+05 | 1.20E+05 | 3.01E+05 | 2.18E+04 | 9.79E+04 | 1000 | 1000 | 60S ribosomal protein L10a OS=Mus musculus GN=Rpl10a PE=1 SV=3 |

| | | | | | | | | | | | |
|---|---|---|---|---|---|---|---|---|---|---|---|
| P60843 | IF4A1 | 5.41 | 4.24E+03 | 4.71E+04 | 3.82E+04 | 1000 | 6.49E+03 | 1000 | 7.86E+03 | 5.83E+03 | Eukaryotic initiation factor 4A-I OS=Mus musculus GN=Eif4a1 PE=1 SV=1 |
| Q8VDQ8 | SIR2 | 5.37 | 1000 | 2.54E+05 | 6.13E+04 | 4.48E+05 | 1.01E+05 | 7.55E+04 | 1000 | 1000 | NAD-dependent protein deacetylase sirtuin-2 OS=Mus musculus GN=Sirt2 PE=1 SV=2 |
| Q6P5H2 | NEST | 5.29 | 7.31E+05 | 6.62E+06 | 2.68E+06 | 8.47E+06 | 6.84E+05 | 2.04E+06 | 7.07E+05 | 1.07E+06 | Nestin OS=Mus musculus GN=Nes PE=1 SV=1 |
| Q3UJU9 | RMD3 | 5.19 | 3.85E+04 | 7.44E+04 | 8.40E+04 | 8.69E+04 | 2.52E+04 | 4.60E+04 | 3.79E+04 | 2.02E+04 | Regulator of microtubule dynamics protein 3 OS=Mus musculus GN=Rmdn3 PE=1 SV=2 |
| Q8C2K5 | RASL3 | 5.17 | 3.45E+04 | 2.84E+04 | 2.56E+04 | 3.80E+04 | 1.73E+04 | 1.62E+04 | 1.18E+04 | 1.10E+04 | RAS protein activator like-3 OS=Mus musculus GN=Rasal3 PE=1 SV=1 |
| P35487 | ODPAT | 5.14 | 3.03E+06 | 7.94E+06 | 4.40E+06 | 2.20E+07 | 9.98E+05 | 5.87E+06 | 9.89E+05 | 1.55E+06 | Pyruvate dehydrogenase E1 component subunit alpha testis-specific form mitochondrial OS=Mus musculus GN=Pdha2 PE=1 SV=1 |
| Q9Z1R2 | BAG6 | 5.08 | 5.03E+06 | 9.02E+06 | 4.03E+06 | 2.02E+07 | 8.84E+05 | 6.52E+06 | 8.06E+05 | 1.66E+06 | Large proline-rich protein BAG6 OS=Mus musculus GN=Bag6 PE=1 SV=1 |
| Q3UHR0 | BAHC1 | 5.06 | 1000 | 1.27E+04 | 2.38E+04 | 2.91E+04 | 1.59E+04 | 1000 | 1000 | 1000 | BAH and coiled-coil domain-containing protein 1 OS=Mus musculus GN=Bahcc1 PE=2 SV=2 |
| Q7SIG6 | ASAP2 | 5.05 | 9.10E+06 | 2.30E+07 | 1.27E+07 | 3.13E+07 | 2.48E+06 | 1.00E+07 | 1.89E+06 | 5.57E+06 | Arf-GAP with SH3 domain ANK repeat and PH domain-containing protein 2 OS=Mus musculus GN=Asap2 PE=1 SV=3 |
| Q6PCX7 | RGMA | 4.99 | 1000 | 6.13E+04 | 4.69E+04 | 3.00E+04 | 1.37E+04 | 1000 | 1.04E+04 | 1.00E+04 | Repulsive guidance molecule A OS=Mus musculus GN=Rgma PE=1 SV=1 |
| Q9CWZ3 | RBM8A | 4.81 | 4.14E+04 | 1.11E+05 | 1.28E+05 | 1.19E+05 | 5.21E+04 | 5.53E+04 | 4.41E+04 | 2.34E+04 | RNA-binding protein 8A OS=Mus musculus GN=Rbm8a PE=1 SV=3 |
| P61028 | RAB8B | 4.8 | 9.62E+03 | 2.15E+05 | 5.55E+04 | 4.86E+04 | 4.39E+04 | 2.35E+04 | 1.21E+04 | 5.49E+03 | Ras-related protein Rab-8B OS=Mus musculus GN=Rab8b PE=1 SV=1 |
| Q0P5X1 | LRIQ1 | 4.69 | 3.28E+05 | 1.35E+06 | 3.68E+05 | 1.18E+06 | 1.18E+05 | 3.17E+05 | 9.25E+04 | 3.45E+05 | Leucine-rich repeat and IQ domain-containing protein 1 OS=Mus musculus GN=Lrriq1 PE=2 SV=2 |
| P52480 | KPYM | 4.63 | 1000 | 4.02E+05 | 3.13E+05 | 8.78E+05 | 4.87E+04 | 3.36E+05 | 5.20E+04 | 1.06E+05 | Pyruvate kinase PKM OS=Mus musculus GN=Pkm PE=1 SV=4 |
| P17156 | HSP72 | 4.52 | 1000 | 1.14E+05 | 1.34E+05 | 1.32E+04 | 3.05E+04 | 2.50E+04 | 1.64E+04 | 1000 | Heat shock-related 70 kDa protein 2 OS=Mus musculus GN=Hspa2 PE=1 SV=2 |
| Q61122 | NAB1 | 4.34 | 5.76E+04 | 5.59E+04 | 9.75E+04 | 3.55E+04 | 2.31E+04 | 3.38E+04 | 3.14E+04 | 1.68E+04 | NGFI-A-binding protein 1 OS=Mus musculus GN=Nab1 PE=1 SV=2 |
| Q61290 | CAC1E | 4.27 | 2.32E+06 | 8.26E+06 | 4.16E+06 | 1.38E+07 | 1.80E+06 | 3.27E+06 | 8.35E+05 | 2.78E+06 | Voltage-dependent R-type calcium channel subunit alpha-1E OS=Mus musculus GN=Cacna1e PE=1 SV=1 |
| P62751 | RL23A | 4.15 | 1.02E+05 | 2.33E+05 | 2.88E+05 | 1.46E+05 | 9.91E+04 | 1.12E+05 | 1.04E+05 | 3.74E+04 | 60S ribosomal protein L23a OS=Mus musculus GN=Rpl23a PE=1 SV=1 |
| Q9QXS1 | PLEC | 4.07 | 1.71E+05 | 1.59E+05 | 1.15E+05 | 1000 | 1000 | 1.21E+05 | 1000 | 1.42E+04 | Plectin OS=Mus musculus GN=Plec PE=1 SV=3 |
| O35594 | IFT81 | 4.02 | 4.30E+06 | 1.05E+07 | 6.01E+06 | 1.68E+07 | 1.52E+06 | 6.83E+06 | 1.21E+06 | 2.49E+06 | Intraflagellar transport protein 81 homolog OS=Mus musculus GN=Ift81 PE=1 SV=4 |
| P47911 | RL6 | 3.99 | 5.52E+04 | 1.02E+05 | 7.08E+04 | 9.85E+04 | 4.10E+04 | 5.04E+04 | 4.91E+04 | 2.12E+04 | 60S ribosomal protein L6 OS=Mus musculus GN=Rpl6 PE=1 SV=3 |
| Q2VIS4 | FILA2 | 3.98 | 4.32E+04 | 1000 | 4.88E+04 | 8.61E+04 | 1000 | 4.32E+04 | 1000 | 1.18E+04 | Filaggrin-2 OS=Mus musculus GN=Flg2 PE=1 SV=2 |
| Q9ES34 | UBE3B | 3.91 | 3.20E+04 | 4.49E+05 | 4.14E+05 | 9.11E+04 | 5.06E+04 | 9.41E+04 | 9.06E+04 | 7.80E+04 | Ubiquitin-protein ligase E3B OS=Mus musculus GN=Ube3b PE=1 SV=3 |
| P04187 | GRAB | 3.73 | 1000 | 4.67E+04 | 1.16E+05 | 2.86E+04 | 1000 | 3.37E+04 | 2.35E+04 | 3.15E+04 | Granzyme B(G H) OS=Mus musculus GN=Gzmb PE=1 SV=1 |
| P67984 | RL22 | 3.67 | 8.64E+04 | 1.51E+05 | 9.59E+04 | 5.69E+04 | 3.88E+04 | 3.22E+04 | 3.72E+04 | 2.26E+04 | 60S ribosomal protein L22 OS=Mus musculus GN=Rpl22 PE=1 SV=2 |
| Q61171 | PRDX2 | 3.63 | 1.26E+05 | 2.96E+05 | 1.26E+05 | 1.52E+05 | 3.12E+04 | 1.11E+05 | 4.90E+04 | 4.60E+04 | Peroxiredoxin-2 OS=Mus musculus GN=Prdx2 PE=1 SV=3 |
| Q9ESD7 | DYSF | 3.57 | 2.28E+05 | 1.63E+05 | 1.79E+05 | 5.74E+04 | 3.14E+04 | 1.87E+04 | 1.46E+05 | 1.94E+04 | Dysferlin OS=Mus musculus GN=Dysf PE=1 SV=3 |
| P99027 | RLA2 | 3.5 | 8.30E+04 | 8.28E+04 | 1.99E+05 | 8.16E+04 | 3.93E+04 | 3.93E+04 | 5.20E+04 | 2.50E+04 | 60S acidic ribosomal protein P2 OS=Mus musculus GN=Rplp2 PE=1 SV=3 |
| Q8K348 | ACV1C | 3.35 | 1000 | 5.48E+04 | 5.15E+04 | 4.61E+04 | 1.68E+04 | 1.62E+04 | 1.41E+04 | 7.72E+03 | Activin receptor type-1C OS=Mus musculus GN=Acvr1c PE=2 SV=3 |
| P62315 | SMD1 | 3.31 | 5.77E+04 | 2.66E+04 | 6.07E+04 | 3.59E+04 | 1.91E+04 | 2.64E+04 | 1.55E+04 | 2.85E+04 | Small nuclear ribonucleoprotein Sm D1 OS=Mus musculus GN=Snrpd1 PE=1 SV=1 |
| B1AY13 | UBP24 | 3.21 | 1000 | 3.11E+05 | 1.99E+05 | 1.27E+05 | 2.36E+04 | 8.02E+04 | 1000 | 1.35E+05 | Ubiquitin carboxyl-terminal hydrolase 24 OS=Mus musculus GN=Usp24 PE=1 SV=1 |
| P14873 | MAP1B | 3.21 | 2.65E+05 | 1.87E+05 | 2.17E+05 | 2.31E+05 | 6.58E+04 | 9.26E+04 | 7.62E+04 | 1.04E+05 | Microtubule-associated protein 1B OS=Mus musculus GN=Map1b PE=1 SV=2 |
| Q8BG05 | ROA3 | 3.2 | 1.21E+05 | 1.30E+05 | 1.75E+05 | 2.10E+05 | 6.18E+04 | 10000 | 6.94E+04 | 4.78E+04 | Heterogeneous nuclear ribonucleoprotein A3 OS=Mus musculus GN=Hnrnpa3 PE=1 SV=1 |
| Q9DBR1 | XRN2 | 3.14 | 1.46E+05 | 6.50E+04 | 1.35E+05 | 7.85E+04 | 2.82E+04 | 5.77E+04 | 3.92E+04 | 3.67E+04 | 5'-3' exoribonuclease 2 OS=Mus musculus GN=Xrn2 PE=1 SV=1 |
| Q8QZY9 | SF3B4 | 3.11 | 5.11E+04 | 2.69E+04 | 10000 | 2.14E+04 | 2.14E+04 | 7.75E+03 | 1.07E+04 | 1.07E+04 | Splicing factor 3B subunit 4 OS=Mus musculus GN=Sf3b4 PE=1 SV=1 |
| Q8VDI1 | ESIP1 | 3.1 | 4.18E+04 | 3.93E+04 | 7.06E+04 | 7.83E+04 | 2.10E+04 | 3.11E+04 | 2.15E+04 | 1.45E+04 | Epithelial-stromal interaction protein 1 OS=Mus musculus GN=Epsti1 PE=1 SV=2 |
| Q9JJN2 | ZFHX4 | 3.07 | 4.59E+04 | 4.11E+04 | 2.85E+04 | 3.45E+04 | 1.46E+04 | 1.35E+04 | 2.10E+04 | 8.75E+03 | Zinc finger homeobox protein 4 OS=Mus musculus GN=Zfhx4 PE=1 SV=1 |
| Q4U4S6 | XIRP2 | 2.93 | 1000 | 1.06E+05 | 2.89E+04 | 4.84E+04 | 1.59E+04 | 1.38E+04 | 2.47E+04 | 1.88E+04 | Xin actin-binding repeat-containing protein 2 OS=Mus musculus GN=Xirp2 PE=1 SV=1 |
| Q80WS3 | FBLL1 | 2.91 | 5.15E+04 | 5.60E+04 | 6.09E+04 | 9.44E+04 | 2.37E+04 | 3.14E+04 | 2.66E+04 | 2.44E+04 | rRNA/tRNA 2'-O-methyltransferase fibrillarin-like protein 1 OS=Mus musculus GN=Fbll1 PE=2 SV=1 |
| Q99N13 | HDAC9 | 2.78 | 3.03E+05 | 5.76E+05 | 3.77E+05 | 5.37E+05 | 1.89E+05 | 1.41E+05 | 1.84E+05 | 2.41E+05 | Histone deacetylase 9 OS=Mus musculus GN=Hdac9 PE=1 SV=2 |
| Q9CPR4 | RL17 | 2.67 | 1000 | 9.82E+04 | 1.33E+05 | 1.01E+05 | 4.74E+04 | 1000 | 5.92E+04 | 3.54E+04 | 60S ribosomal protein L17 OS=Mus musculus GN=Rpl17 PE=1 SV=3 |
| P30658 | CBX2 | 2.66 | 1.30E+06 | 3.57E+06 | 3.00E+06 | 2.89E+06 | 1.27E+06 | 1.68E+06 | 1.34E+06 | 4.46E+05 | Chromobox protein homolog 2 OS=Mus musculus GN=Cbx2 PE=1 SV=2 |
| A2AAJ9 | OBSCN | 2.65 | 1.25E+05 | 6.31E+04 | 3.96E+04 | 3.66E+04 | 2.00E+04 | 2.33E+04 | 5.03E+04 | 2.02E+04 | Obscurin OS=Mus musculus GN=Obscn PE=1 SV=2 |
| Q62036 | CP131 | 2.62 | 1000 | 1.40E+06 | 5.74E+05 | 2.45E+06 | 1.04E+05 | 6.02E+05 | 1.06E+06 | 1.73E+05 | Centrosomal protein of 131 kDa OS=Mus musculus GN=Cep131 PE=1 SV=2 |
| Q8VCH8 | UBXN4 | 2.58 | 7.84E+04 | 1.96E+05 | 1.33E+05 | 1.09E+05 | 8.21E+04 | 4.89E+04 | 6.01E+04 | 3.63E+04 | UBX domain-containing protein 4 OS=Mus musculus GN=Ubxn4 PE=1 SV=1 |
| P42932 | TCPQ | 2.57 | 1.54E+04 | 3.79E+04 | 4.87E+04 | 5.78E+04 | 1.64E+04 | 2.63E+04 | 1.74E+04 | 9.83E+03 | T-complex protein 1 subunit theta OS=Mus musculus GN=Cct8 PE=1 SV=3 |
| Q8VIJ6 | SFPQ | 2.53 | 1000 | 2.87E+04 | 2.18E+04 | 3.00E+04 | 4.60E+03 | 2.96E+04 | 1000 | 5.45E+03 | Splicing factor proline- and glutamine-rich OS=Mus musculus GN=Sfpq PE=1 SV=1 |
| Q9Z319 | CORIN | 2.53 | 1.67E+05 | 6.14E+05 | 2.18E+05 | 6.16E+05 | 1.55E+05 | 8.93E+04 | 2.65E+05 | 2.15E+05 | Atrial natriuretic peptide-converting enzyme OS=Mus musculus GN=Corin PE=2 SV=2 |
| Q91VW5 | GOGA4 | 2.5 | - | 1.88E+05 | 4.85E+04 | 5.26E+05 | 1.50E+05 | 2.05E+05 | 1.56E+04 | 8.52E+04 | Golgin subfamily A member 4 OS=Mus musculus GN=Golga4 PE=1 SV=2 |
| Q8CCS6 | PABP2 | 2.49 | 1.71E+05 | 1.01E+05 | 2.04E+05 | 1.48E+05 | 7.60E+04 | 7.63E+04 | 7.58E+04 | 5.39E+04 | Polyadenylate-binding protein 2 OS=Mus musculus GN=Pabpn1 PE=1 SV=3 |
| Q99M28 | RNPS1 | 2.48 | 1.52E+05 | 3.42E+05 | 1.78E+05 | 2.80E+05 | 1.16E+05 | 1.14E+05 | 1.41E+05 | 6.07E+04 | RNA-binding protein with serine-rich domain 1 OS=Mus musculus GN=Rnps1 PE=1 SV=1 |
| Q61768 | KINH | 2.45 | 2.69E+04 | 3.96E+04 | 2.14E+04 | 10000 | 1.51E+04 | 1.54E+04 | 1.14E+04 | 1.10E+04 | Kinesin-1 heavy chain OS=Mus musculus GN=Kif5b PE=1 SV=3 |
| Q61990 | PCBP2 | 2.43 | 1000 | 6.68E+04 | 5.55E+04 | 3.04E+04 | 2.37E+04 | 1.51E+04 | 1.72E+04 | 1.36E+04 | Poly(rC)-binding protein 2 OS=Mus musculus GN=Pcbp2 PE=1 SV=1 |
| Q8BP47 | SYNC | 2.39 | 2.30E+05 | 1.87E+05 | 8.06E+04 | 3.02E+05 | 8.74E+04 | 1.22E+05 | 5.95E+04 | 1.03E+05 | Asparagine--tRNA ligase cytoplasmic OS=Mus musculus GN=Nars PE=1 SV=2 |
| Q6PB66 | LPPRC | 2.37 | 2.66E+04 | 4.87E+04 | 2.63E+04 | 3.48E+04 | 1.41E+04 | 1.89E+04 | 1.90E+04 | 1.10E+04 | Leucine-rich PPR motif-containing protein mitochondrial OS=Mus musculus GN=Lrpprc PE=1 SV=2 |
| Q8K3K7 | PLCB | 2.34 | 2.76E+06 | 4.54E+06 | 1.98E+06 | 2.20E+06 | 1.52E+06 | 1.04E+06 | 1.94E+06 | 9.77E+05 | 1-acyl-sn-glycerol-3-phosphate acyltransferase beta OS=Mus musculus GN=Agpat2 PE=1 SV=1 |
| Q8BX22 | SALL4 | 2.33 | 1.98E+05 | 10000 | 3.22E+05 | 2.72E+05 | 9.38E+04 | 1.59E+05 | 8.08E+04 | 1.65E+05 | Sal-like protein 4 OS=Mus musculus GN=Sall4 PE=1 SV=2 |
| P61327 | MGN | 2.33 | 2.94E+05 | 3.97E+05 | 4.45E+05 | 5.83E+05 | 2.02E+05 | 2.37E+05 | 2.41E+05 | 1.33E+05 | Protein mago nashi homolog OS=Mus musculus GN=Magoh PE=2 SV=1 |
| Q9Z1K7 | APC2 | 2.31 | 8.20E+04 | 1.22E+05 | 5.90E+04 | 8.28E+04 | 4.48E+04 | 3.86E+04 | 3.87E+04 | 10000 | Adenomatous polyposis coli protein 2 OS=Mus musculus GN=Apc2 PE=1 SV=1 |
| O70133 | DHX9 | 2.27 | 7.52E+04 | 7.45E+04 | 8.57E+04 | 6.79E+04 | 3.59E+04 | 4.35E+04 | 3.85E+04 | 2.70E+04 | ATP-dependent RNA helicase A OS=Mus musculus GN=Dhx9 PE=1 SV=2 |
| Q9QXK7 | CPSF3 | 2.26 | 1000 | 1.22E+05 | 8.19E+04 | 4.24E+04 | 2.54E+04 | 3.78E+04 | 3.04E+04 | 2.37E+04 | Cleavage and polyadenylation specificity factor subunit 3 OS=Mus musculus GN=Cpsf3 PE=1 SV=2 |
| Q9D8E6 | RL4 | 2.23 | 7.14E+04 | 7.60E+04 | 1.18E+05 | 1.04E+05 | 4.94E+04 | 6.83E+04 | 4.19E+04 | 1.92E+04 | 60S ribosomal protein L4 OS=Mus musculus GN=Rpl4 PE=1 SV=3 |
| Q33DR3 | DLP1 | 2.21 | 5.52E+04 | 1.49E+05 | 1.16E+05 | 8.84E+04 | 5.88E+04 | 6.36E+04 | 5.91E+04 | 1.71E+04 | Decaprenyl-diphosphate synthase subunit 2 OS=Mus musculus GN=Pdss2 PE=1 SV=2 |
| P51410 | RL9 | 2.19 | 9.55E+04 | 9.19E+04 | 1.21E+05 | 6.05E+04 | 3.47E+04 | 5.75E+04 | 5.02E+04 | 3.72E+04 | 60S ribosomal protein L9 OS=Mus musculus GN=Rpl9 PE=2 SV=2 |
| Q68FF6 | GIT1 | 2.15 | 2.10E+04 | 10000 | 5.38E+04 | 1.44E+05 | 4.41E+04 | 5.52E+04 | 3.19E+04 | 1.24E+04 | ARF GTPase-activating protein GIT1 OS=Mus musculus GN=Git1 PE=1 SV=1 |

**Supplemental Table 1**

**Supplemental Table 1. Mass spec characterization of proteins immunoprecipitated from day 7 GVHD T cells using a phospho-AMPK motif specific antibody** WT versus AMPK KO T cells were transplanted into B6D2F1 recipients, recovered on day 7, and cell lysates immunoprecipitated using an antibody recognizing the phosphorylated AMPK-specific motif LxRxx(pS/pT). Precipitated proteins were subsequently processed via LC-MS to identify candidates expressed at 3-fold or higher levels in WT compared to AMPK KO T cells.

**Supplemental Table 2**

| *Flow cytometry antibodies* | | | | |
|---|---|---|---|---|
| *Antigen* | *Conjugate* | *Company* | *Clone* | *Catalog Number* |
| *Human CD3* | *PE* | *BioLegend* | *UCHT1* | *300408* |
| *Human CD45* | *PerCPCy5.5* | *BioLegend* | *HI30* | *304207* |
| *CD3* | *PerCPCy5.5* | *eBioscience* | *145-2C11* | *45-00310-80* |
| *CD4* | *BUV737* | *BD Biosciences* | *GK1.5* | *612761* |
| *CD4* | *FITC* | *eBioscience* | *RM4.4* | *11-0043-85* |
| *CD8* | *eFllur780* | *eBioscience* | *53-6.7* | *47-0081-82* |
| *CD45.1* | *BV711* | *BioLegend* | *A20* | *110739* |
| *CD90.1* | *PerCPCy5.5* | *eBioscience* | *HIS51* | *45-0900-82* |
| *CD90.2* | *FITC* | *BD Biosciences* | *30-H12* | *553012* |
| *IFNγ* | *PE* | *eBioscience* | *XMG1.2* | *12-7311-82* |
| *IFNγ* | *APC* | *eBioscience* | *XMG1.2* | *17-7311-81* |
| *IL-17a* | *PECy7* | *eBioscience* | *eBio17B7* | *25-7177-80* |
| *TCR-β* | *PerCPCy5.5* | *eBioscience* | *H57-597* | *45-5961-82* |
| *TNF* | *APC* | *eBioscience* | *MD6-XT22* | *17-7321-81* |

Supplementary Table 3 – Reagents for flow cytometry

| *Additional flow cytometry reagents* | | | |
|---|---|---|---|
| *Reagent* | *Ex/Em* | *Company* | *Catalog Number* |
| *CellTrace Violet* | *405/450* | *Invitrogen* | *C34557* |
| *Live/Dead Aqua* | *375/526* | *Invitrogen* | *L34965* |

Supplementary Table 4 – Antibodies for Immunoblotting

| *Antigen* | *Company* | *Catalog number* | *Clone* |
|---|---|---|---|
| *Aldolase A* | *Cell Signaling* | *8060* | *D73H4* |
| *B-actin* | *Cell Signaling* | *4970* | *13E5* |

| | | | |
|---|---|---|---|
| *GAPDH* | *Cell Signaling* | *5174* | *D16H11* |
| *Ulk-1* | *Cell Signaling* | *8054* | *D8H5* |
| *Phospho-ULK-1 (ser555)* | *Cell Signaling* | *5869* | *D1H4* |

Abbreviations: GAPDH – glyceraldehyde 3-phosphate dehydrogenase , ULK-1 – Unc51-like kinase 1.
